# The morphology and small subunit rDNA gene phylogeny of the novel goniomonad genus *Ebisugoniomonas* and two novel *Poseidogoniomonas* species

**DOI:** 10.1101/2025.02.23.639744

**Authors:** Daryna Zavadska, Maria Sachs, Hartmut Arndt, Daniel J. Richter

## Abstract

Goniomonads are heterotrophic representatives of Cryptista, sister lineage to the phototrophic Cryptists. Goniomonads are morphologically and genetically diverse, comprising nine genera described so far. Besides morphologically described genera, SSU rDNA sequencing data has revealed the existence of at least one more monophyletic genus-level lineage within goniomonads, as well as more novel species within the described genera. Furthermore, the diversity of SSU rDNA sequences within goniomonads is not fully investigated, and addition of novel sequences can be crucial for understanding the evolution and diversity of goniomonads and for providing a basis for taxonomic descriptions. The morphological data record available for goniomonads is relatively limited: many cellular features remain undescribed, or described only for a few isolates.

In this study, we describe the new genus and species *Ebisugoniomonas hispanica* n. gen. n. sp., and two novel species within *Poseidogoniomonas*: *P. posteriocircus* n. sp. and *P. zopfkaesiformis* n. sp. We report a range of observations on the cellular features of goniomonads: periplast plate arrangement pattern and intracellular structures. We also update the SSU rDNA phylogeny of goniomonads by adding sequences from novel isolates and environmental DNA studies.

Several sequences recovered from novel isolates and environmental DNA datasets likely represent highly divergent lineages of goniomonads in the SSU rDNA phylogeny.

## 1 Introduction

Goniomonads are a monophyletic lineage within Cryptista. Morphologically, all goniomonads studied so far are aplastidic, biflagellated, laterally flattened cells, with a truncate anterior and a conspicuous horizontal band of ejectosomes on the anterior side of the cell (Kim and Archibald, 2013; Larsen and Patterson, 1990; Novarino, 2003; Patterson and Larsen, 1992; Sachs et al., 2025; Schuster, 1968).

The phylogenetic position of goniomonads within Cryptista, their cell biology, and their genomic sequences have been of interest primarily because goniomonads are heterotrophic relatives of cryptomonads, the photosynthetic cryptists. Here we use “cryptomonads” to refer to Cryptomonadales *sensu* Hoef-Emden and Archibald (2017), (Adl et al., 2019), or Cryptophyceae *sensu* Von der Heyden and Cavalier-Smith (2004), (Clay, 2015)). Therefore, investigating goniomonads was appraised to be particularly important for understanding the evolution of photosymbioses in Cryptista as they are considered as a “pre-secondary endosymbiosis cryptomonad” (McFadden, 2018).

According to the multiple phylogenetic trees reconstructed throughout the past three decades based on SSU rDNA gene sequences (Deane et al., 2002; Marin et al., 1998; Martin-Cereceda et al., 2010; McFadden and Hill, 1994), phylogenomic trees inferred from the alignment of 157 protein sequences (Yabuki et al., 2014) and 250 protein sequences (Cenci et al., 2018), goniomonads branch as a sister lineage to the plastid-containing cryptomonads. With the current placement of goniomonads within Cryptista, a scenario of a single plastid gain in the cryptophyte common ancestor (McFadden and Hill, 1994; Tanifuji and Onodera, 2017; Yabuki et al., 2014) is more parsimonious compared to the scenario of multiple plastid loss in the ancestor of heterotrophic lineages of Cryptista (the latter scenario aligns with chromalveolate hypothesis (Cavalier-Smith, 1999; Von der Heyden and Cavalier-Smith, 2004)). No significant evidence of genes transferred from a secondary endosymbiotic plastid was found based on genomic studies (Cenci et al., 2018). Finally, no ultrastructural evidence for plastid-like compartments was found (Cenci et al., 2018).

Although the position of goniomonads as a monophyletic lineage of heterotrophic cryptists sister to cryptomonads is well-resolved and unambiguous, diversity and divergence within the lineage is profound. Yet, until recently, it remained largely unexplored.

The deep genetic divergence between freshwater and marine goniomonads, as well as the high divergence of SSU rDNA sequences within both freshwater and marine clades, was established two decades ago (Martin-Cereceda et al., 2010; Von der Heyden and Cavalier-Smith, 2004). However, until recently, all the representatives of goniomonads were assigned to a single genus Goniomonas Stein 1878, with only five species formally described (Sachs et al., 2025). Taxonomic revision of *Goniomonas* Stein 1878 included splitting the genus into nine genera: *Limnogoniomonas, Aquagoniomonas* and *Goniomonas* for freshwater lineages, and *Marigoniomonas, Thalassogoniomonas, Poseidogoniomonas, Cosmogoniomonas, Baltigoniomonas* and *Neptunogoniomonas* for marine lineages, based on the pairwise distances and synapomophy analysis between partial SSU rDNA sequences and, in most cases, on morphological differences between type strains.

In the SSU rDNA phylogeny of goniomonads, which was used as a basis for their taxonomic revision, a distinct sequence cluster appears as sister to four described marine genera, branching between them and two genera with known brackish-water and marine representatives: *Baltigoniomonas* and *Neptunogoniomonas*. This cluster clearly represents another divergent lineage of goniomonads, for which we have isolates allowing formal description as a separate genus. Furthermore, within the *Poseidogoniomonas* genus only a single species has been described thus far, although more isolates from this lineage have been sequenced and await morphological description. In this study, we describe one species of the novel early-branching goniomonad genus, *Ebisugoniomonas hispanica* n. gen. n. sp., and two novel species of *Poseidogoniomonas*: *Poseidogoniomonas posteriocircus* n. sp. and *Poseidogoniomonas zopfkaesiformis* n. sp.

Finally, the knowledge on the diversity of cryptists was recently expanded by environmental sequencing data, which uncovered the existence of several lineages, including CRY-1 (Piwosz et al., 2016), CRY-2 (Shalchian-Tabrizi et al., 2008) and CRY-3 (Yabuki et al., 2014). Together with the fast accumulation of novel goniomonad SSU rDNA sequences, it is reasonable to assume that the substantial diversity of goniomonads remains uncultured, likely available as environmental sequencing data. In this study, we expand the SSU rDNA of goniomonad sequence dataset with environmental sequencing data, and use it for phylogenetic tree inference. New sequences incorporated into the phylogeny in this study reveal the presence of even more novel goniomonad lineages, as well as additional diversity within the previously established clades.

In terms of goniomonad morphology and function, the cells of goniomonads have a symmetry different from that observed in cryptomonads. Goniomonads are flattened in a way such that the cell is more narrow when observed from the dorsal (where flagella are located (Clay, 2015; Hoef-Emden and Archibald, 2017)) or ventral (where the groove extends) sides, and wide and flat in lateral projection (Von der Heyden and Cavalier-Smith, 2004). Note that in this study we use the definition of dorsal and ventral sides according to Mignot (1965), defining the wide flat sides as “left” and “right", based on their position when observed from the dorsal side. Correspondingly to the symmetry, the movement of goniomonads also differs from that of typical cryptomonads (Hoef-Emden and Archibald, 2017; Novarino, 2003) and its most common form of movement has been described as “jerky” or irregular gliding along a substrate (Kim and Archibald, 2013; Martin-Cereceda et al., 2010) with one of the lateral flattened sides facing the substrate (Novarino, 2003). Goniomonads were reported to use oil as storage product (https://www.algaebase.org/search/genus/detail/?genus_id=G479291fbb5b4cb2e). The enzymes for beta-linked polysaccharide metabolism as well as the complete pathway for the production of an alpha glucan storage polysaccharides were putatively identified in the genome and the possibility of *Goniomonas* synthesising beta-storage polysaccharides and starch has been discussed (Cenci et al., 2018).

The morphological features consistently used as diagnostic characters for distinguishing goniomonad species have so far included the number of periplast plates, the length of the flagella (and the presence of mastigonemes) and the ejectosome row and the length as well as width of the cell (Novarino, 2003). Very recently the pattern of periplast plate arrangement was proposed as another diagnostic character in goniomonad taxonomy (Phanprasert et al., 2024), and in this study, we independently confirm and expand this finding. Although some more detailed observations have been made for the surface structure of goniomonads, for the cytoskeletal structure (Kim and Archibald, 2013; Martin-Cereceda et al., 2010), movement patterns and the ejectosome morphology (Kim and Archibald, 2013; Phanprasert et al., 2024; Schuster, 1968), there is a lack of information on how the reported features vary across the breadth of goniomonad diversity. Therefore, in this study, we attempt to gather additional information about the arrangement of periplast plates, movement patterns and intracellular structures of the isolates investigated.

## 2 Materials and Methods

### 2.1 Sampling and cultivation

Plankton samples were obtained from the subsurface water of the Mediterranean Sea near Blanes Bay, Spain, 41°40’N 2°40’E throughout the year 2021-2022. Samples were enriched with RS medium (https://mediadive.dsmz.de/medium/P4, https://mediadive.dsmz.de/medium/P5) of nutrient (NM) to non-nutrient (NNM) component ratios between 1:5 and 1:500. Stable mixed cultures were obtained from the samples enriched with the nutrient component of RS medium and sustained in RS medium at +18°C; 4 sequential rounds of dilution to extinction were performed independently on each stable mixed culture to obtain monoclonal isolates of goniomonads.

The isolates we used for the species description were monoclonal cultures obtained from initial stable mixed cultures: the monoclonal isolate BEAP0340 was obtained from the mixed culture of BEAP0075; monoclonal BEAP0338 was obtained from the mixed culture of BEAP0079; monoclonal BEAP0335 was obtained from the mixed culture of BEAP0220; monoclonal BEAP0342 was obtained from the mixed culture of BEAP0128. SSU rDNA sequences were obtained as described below from **both** goniomonads in the initial mixed cultures and for the monoclonal strains derived from these cultures by the dilution to extinction. This was done to verify the sequence of monoclonal cultures.

All isolates can grow in RS medium with nutrient to non-nutrient component ratios of 1:50 and 1:100, as well as in artificial seawater with an autoclaved rice grain. Monoeukaryotic clonal strains are deposited in the Roscoff Culture Collection with identifiers 11327 (BEAP0340), 11329 (BEAP0338) and 11333 (BEAP0335).

### 2.2 Morphological studies

Light microscopy images were obtained on a Zeiss Axio Observer.A1 inverted microscope with a water immersion condenser and 100x/1.4NA Oil immersion Differential Interference Contrast (DIC) objective and Allen Video-enhanced system (Hamamatsu C6489, Argus-20).

To prepare the samples for Scanning Electron Microscopy (SEM), cultures were centrifuged at 11000 x g for 8 minutes at room temperature (RT), the pellet was resuspended and fixed in 2.5% v/v glutaraldehyde in PBS for 30 minutes (for BEAP0335 and BEAP0338) or 1 hour (for BEAP0340) at +4°C. Fixed cells were collected on 0.8 µm filters and dehydrated in graded ethanol series (10%, 20%, 30%, 50%, 70%, 80%, 90%, 96%, 96% (overnight) for BEAP0340 and 20%, 30%, 50%, 70% (overnight), 80%, 90%, 96%, 96% for BEAP0335 and BEAP0338) with the first rinse lasting 10 minutes, and each following rinse lasting five minutes more than the previous one.

After dehydration, samples were critical-point dried using liquid carbon dioxide in a Leica EM CPD300 unit (Leica Microsystems, Austria). The dried filters were placed on stubs with colloidal silver and then sputtered with 5 nm gold (for BEAP0335 and BEAP0338) or 5 nm Iridium (for BEAP0340) in a Q150R S (Quorum Technologies, Ltd.) Samples were observed with a Field-Emission Hitachi SU8600 SEM (Hitachi High Technologies Co., Ltd, Japan) at an accelerating voltage of 2 kV or 5 kV.

### 2.3 Molecular studies

To prepare genomic DNA, 100 ml of each monoclonal cell culture was centrifuged at 11000 x g for 8 minutes at RT. Total DNA was extracted using the DNeasy PowerSoil Pro Kit (QIAGEN, cat. no. 47017, Hilden, Germany), according to the manufacturer’s protocol, additionally passing the cell lysate 5 times through a 25 gauge needle after mechanical lysis.

SSU rDNA sequence was amplified using universal eukaryotic 18S primers 18S-42F 5’-CTCAARGAYTAAGCCATGCA-3’ and 18S-1510R 5’-CCTTCYGCAGGTTCACCTAC-3’ (Lopez-Garcia et al., 2003; Vaulot et al., 2022). The PCR protocol was as follows: an initial denaturation step at 94°C for 3 min, followed by 30 cycles at 94°C for 15 s, 58°C for 30 s, 72°C for 3 min and a final extension-terminal A addition step at 72°C for 37 min.

PCR products were visualised on a 1% agarose gel, and products of 1-2kb in length were cloned into the pCR4-TOPO Vector using the TOPO TA cloning kit for sequencing (Invitrogen, product ref. 45-0030, Carlsbad, USA). The ligation product was transfected into homemade Top-10 chemocompetent cells, cells were plated onto LB plates with 100 µg/ml ampicillin and 10 mg/ml X-gal. Blue-white screening allowed for the selection of white bacterial colonies which should have PCR product integrated into the plasmid. White colonies were picked and subjected to colony PCR using M13F and M13R primers. The PCR protocol was as follows: an initial denaturation step at 94°C for 3 min, followed by 30 cycles at 94°C for 20 s, 55°C for 20 s, 72°C for 3 min, and a final extension at 72°C for 5 min. Amplicons of size 1-2 kb were purified from the PCR mix with a Gelpure kit (NZYtech, cat. no MB01101, Lisbon, Portugal), according to the manufacturer’s protocol, except that the DNA was eluted with 30 µl of EcoLav water pre-heated to 40°C. Each amplicon was Sanger-sequenced (Eurofins Genomics, Cologne, Germany) from both ends, using M13F and M13R primers.

Multiple colonies were screened by colony PCR for the transformants with SSU rDNA inserts from each isolate (12 for BEAP0340, 6 for BEAP0335 and 2 for BEAP0338). This was done to verify that the isolates are monoclonal cultures.

The same DNA extraction, PCR, cloning and sequencing protocol was used to produce SSU rDNA sequences for numerous other goniomonad isolates from BEAP lab which are not described in detail in this manuscript because the cultures died shortly after sequencing; nevertheless, these sequences were included in the phylogenetic analysis and were deposited to GenBank under IDs PV090901-PV090922.

### 2.4 Phylogenetic analysis

The dataset obtained from the alignment used in Sachs et al. (2025) was further extended by adding sequences from EukRibo v.2 (Berney et al., 2022): all the sequences classified as “Cryptophyceae” by EukRibo taxonomy were added to the master alignment. The EukRibo database was chosen because (i) the database is manually curated so it is unlikely to contain chimaeras, misclassified sequences, duplicated or highly redundant sequences and (ii) it contains long-read sequencing data from environmental DNA sequencing projects, in particular, the long-read sequences from Jamy et al. (2020).

In addition, we included sequences from the guide alignment kindly provided by Cédric Berney, which contained 240 sequences classified as Cryptista from either EukRibo, NCBI or Jamy et al. (2020) dataset.

The set of sequences from all of the sources listed above was aligned using MAFFT v7.453 (default parameters) (Katoh et al., 2002). The resulting alignment was visualised and manually edited in JalView (Waterhouse et al., 2009). Within goniomononads, only 100% identical sequences were removed.

Finally, to recover sequences which might belong to goniomonads, but were not annotated as such in NCBI nt database, we iteratively searched NCBI nt database using eukref_gbretrieve.py (https://github.com/eukref/pipeline/blob/master/eukref_gbretrieve.py) script from the EukRef pipeline (del Campo et al., 2018) (a congruous approach for retrieving environmental sequences was used in Flegontova et al. (2018) and subsequently in Packer et al. (2025)). All sequences of goniomonads from the dataset described above were clustered and sorted with usearch v8.0.1517 (Vardanian, 2023), and used as input for the iterative search against NCBI nt; the GenBank BLAST cutoff (“–idnt”) for the iterative search was set to 80, while the most inclusive PR2 taxonomic group name was “Goniomonadales".

Two sequences, previously not included in the dataset, were recovered by this search: EF526920 and EF674349. The two sequences were added to the dataset created in the previous step; the resulting dataset was aligned with MAFFT version 7.453 (Katoh et al., 2002) using default parameters. FastTree version 2.1.11 (Price et al., 2010) with default settings was used to predict the approximate position of the EF526920 and EF674349 on the phylogeny of Cryptista and to define the outgroup for the final alignment. With EF526920 placed within CRY-4 lineage and EF674349 placed within goniomonads, we added these sequences into the dataset as well. From all the non-goniomonad sequences, nine SSU rDNA sequences of CRY-1 lineage were selected as an outgroup for the phylogeny presented in this study (Figures 5, S4). In addition, phylogenetic trees with a larger outgroup were run, to (i) check more precisely positions of EF674349 and EF526920 and (ii) check how the general topology of the tree is altered when more diverse outgroup is taken. Those trees are presented in FiguresS5 and S6.

The resulting datasets included sequences from all the aforementioned sources combined, with either CRY-1 or all Cryptista sequences as an outgroup. Duplicated sequences were removed using the -rdump function from the seqkit tool both before running MAFFT alignment, and after trimming (Shen et al., 2016).

Sequences were aligned with MAFFT v7.453 (Katoh et al., 2002) using the L-INS-I algorithm, with a maximum number of iterations set to 1000. The resulting alignment was trimmed with trimAl (Capella-Gutiérrez et al., 2009) using several options: -gappyout option, and -gt option with gap trimming cutoffs of 0.1, 0.2 and 0.5. The trimmed alignments were assessed by estimating the number and percentage of parsimony-informative sites with the corresponding function from phykit (Steenwyk et al., 2021), and visually inspected. We found that the optimal trimming setting was trimAl with the -gt option with a threshold of 0.2.

The final alignment used in phylogeny reconstruction contained 97 sequences. ModelTest-NG v0.2.0 (Darriba et al., 2019) was run on the trimmed alignment; GTR+I+G4 was identified as the most optimal model, according to AIC and AICc and it was used for both Bayesian (BI) and Maximum Likelihood (ML) analysis described below (HKY+I+G4 model was the most optimal according to BIC but due to the fact that not all versions of RAxML support this model, for the full comparability of BI and ML phylogenies, GTR+I+G4 model was used for both methods). ML phylogeny was inferred using RAxML version 8.2.12, with 1000 bootstrap replicates. Bayesian phylogeny was inferred using MrBayes v3.2.6 (Ronquist and Huelsenbeck, 2003), with sampling frequency 10 and stop value of topological convergence set as 0.01. With an initial number of generations set to 1,000,000, the average standard deviation of split frequencies reached a plateau between 0.09-0.012 between generations 600,000-1,000,000. The run was stopped at generation 1,000,000 with the average standard deviation of split frequencies being 0.01.

Pairwise cophenetic distances were calculated from both ML and Bayesian trees with the approach described in Richter et al. (2018), with the ape R package v.5.7-1.

### 2.5 Data visualisation, postprocessing and availability

Morphological measurements were conducted using ZEN (blue edition) imaging software v.3.5.093.00001 or v.3.6.095.04000, and visualised using a custom R script. Images for the publication were prepared using Fiji (Schindelin et al., 2012) and Inkscape. Imaging data associated with the descriptions can be accessed on Figshare: https://figshare.com/s/173e3e1de30fd1151fc8.

Branches of the final trees were first renamed automatically using a custom R script. The resulting trees with renamed leaves were visualised in TV-bot (Xie et al., 2023), and finally, edited in Inkscape. Intermediate files obtained in the process of inferring phylogenetic trees and the input sequence dataset, along with the sequence of commands used are available on GitHub: https://github.com/beaplab/Goniomonas.git.

SSU rDNA sequences of the BEAP isolates used in this study and obtained in the Biology and Ecology of Abundant Protist lab are deposited in GenBank under IDs PV090901-PV090922. The list of GenBank IDs corresponding to SSU rDNA of BEAP isolates from this study is available by the link https://github.com/beaplab/Goniomonas/blob/main/GENBANK_ids_BEAP_isolates.tsv.

## 3 Results

### 3.1 *Ebisugoniomonas* n. gen. isolates

#### 3.1.1 BEAP0340

The cells are colourless heterotrophic biflagellates of 4.8-6.4 µm long and 4.3-5.6 µm wide; two anterio-laterally oriented flagella are of an unequal length, with the anterior flagellum of 5.5-7.5 µm in length and the posterior flagellum of 8.3-10.2 µm in length (Figure 1). With the flagella of unequal length, the most frequent movement pattern observed in this strain involved only the anterior flagellum beating actively, while the posterior flagellum is mainly observed trailing, producing occasional weak strokes (Suppl. Material S1).

**Figure 1.**
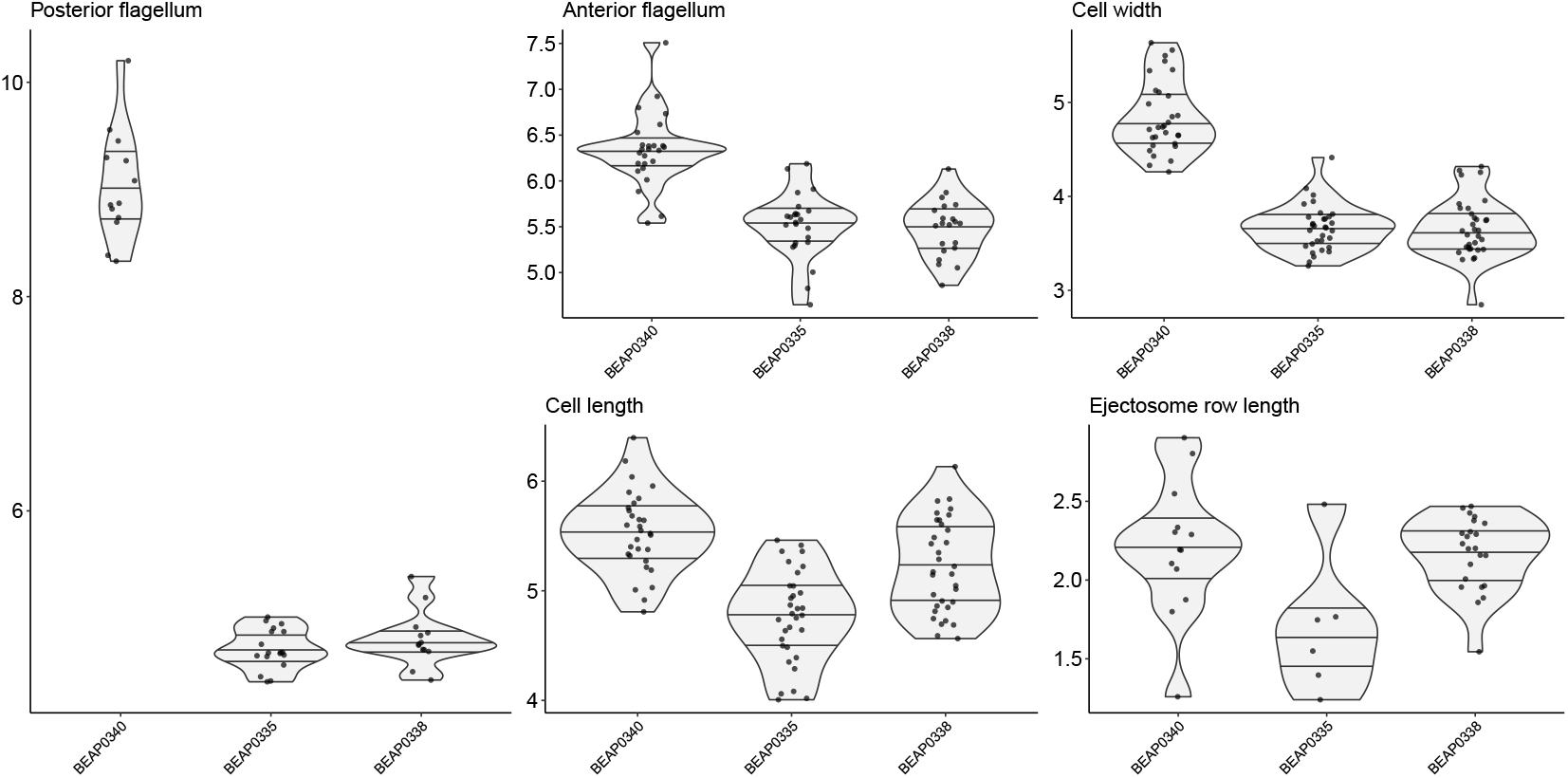
Summary of morphological parameters measured for the isolates described. Each violin plot illustrates the distribution of the given parameter measured for a given isolatespecies (BEAP0335 - *Poseidogoniomonas zopfkaesi-formis* n. sp., BEAP0338 - *Poseidogoniomonas posteriocircus* n. sp. and BEAP0340 - *Ebisugoniomonas hispanica* n. gen. et n. sp.); the lines on the violin plots represent the quartiles. Each dot corresponds to a measurement obtained from one cell. All the values presented are in µm; note the different vertical axis scales.

The cells are laterally compressed and close to roundish shape, with width almost equal to their length. The band of ejectosomes is 1.3-2.9 µm long (Figure 1), and was observed in roughly half of the adult cells analysed. Another interesting feature consistently observed for this isolate during DIC microscopy is the presence of two, or, less frequently, one toroid-shaped structure below the anterior extension, close to the row of ejectosomes (Figure 2-B). Those structures are clearly different from ejectosomes themselves, and to our current knowledge have not been reported in any other described goniomonad. The cytoplasm of fully-grown cells (just before fission), and especially of the old cells, is relatively heavily vacuolised (highly likely, lipid globules; Figure 2-B, Figure S2).

**Figure 2.**
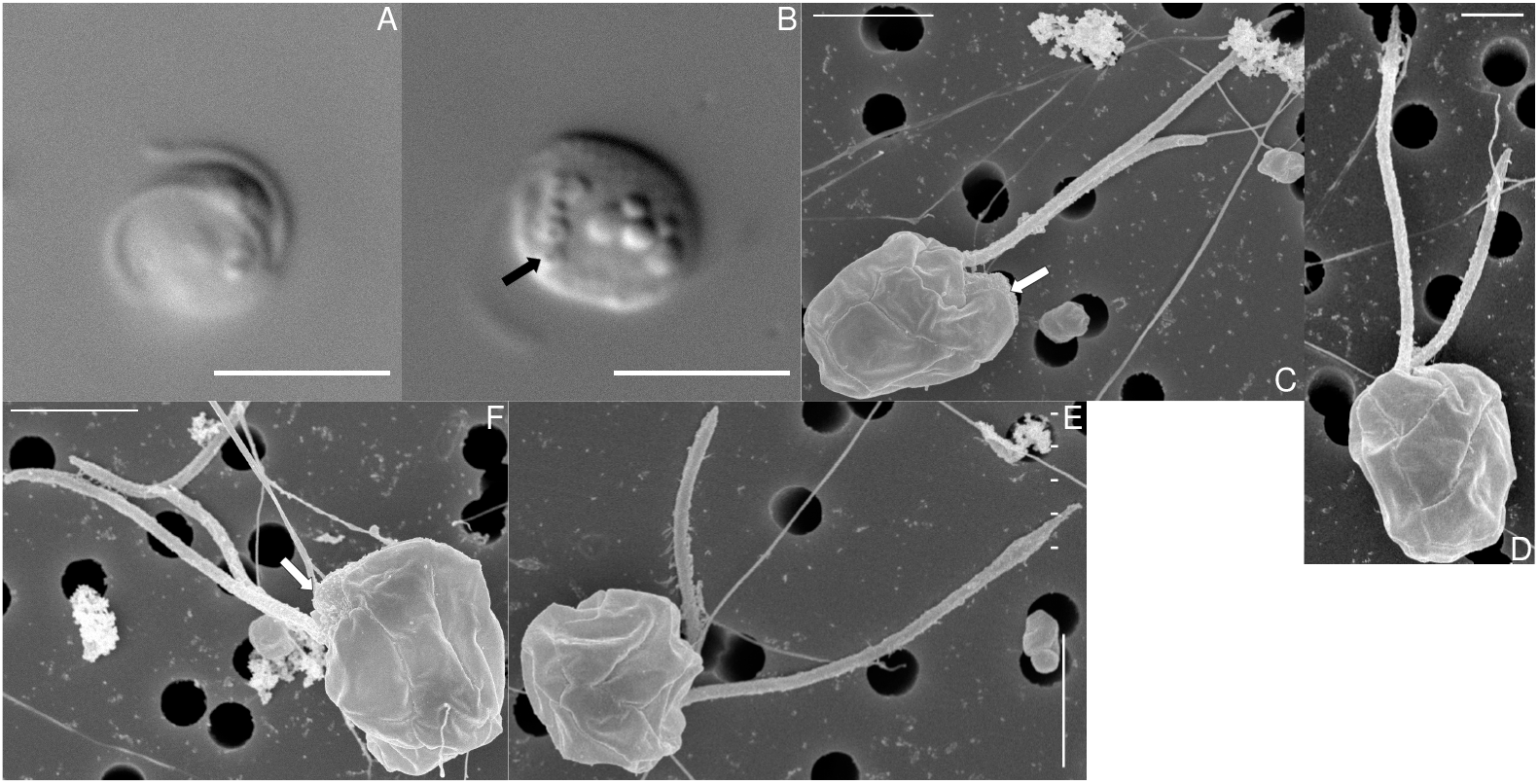
Light and electron micrographs showing *Ebisugoniomonas* n. gen. isolate described in this study. *Ebisugoniomonas hispanica* n. gen. et n. sp.; bar indicates 1 µm (D), 2 µm (C, E-F) and 5 µm (A-B). Arrows indicate a toroid-shaped structure (B), and a structure resembling a granular zone (C, F).

In BEAP0340 five periplast plates (on the right and left sides) are present. The two plates located in front of the anterior depression form a conspicuous extension of the anterior part of the cell; they are oriented at an acute angle to the anterior end. The upper plate (the one closer to the anterior part) joins with the triangular plate located at the anterior depression. On its other side, the triangular plate is adjacent to the elongated plate which itself joins all the other periplast plates (Figures 2-C-F, 4-A). Note that the elongated plate can appear to be split in two parts in some cells -the bottom part and the dorsal part. A structure resembling what was described as granular zone in Martin-Cereceda et al. (2010) is present at the anterior end of the cells (Figure 2-F).

The clade with sequences from isolates BEAP0073, BEAP0077, BEAP0078, BEAP0096, BEAP0340 and HQ529498 has 100% Bootstrap support and 1.0 posterior probability. The shortest distances between the *Ebisugoniomonas* n. gen. and *Neptunogoniomonas* or *Baltigoniomonas* representatives is >5%. The maximum pairwise distance observed within the clade is 1.35% (ML) or 1.28% (BI) between HQ529498 and BEAP0340 (Figure 6). Sequence variation within the *Ebisugoniomonas* clade/genus is hard to assess, since the majority of BEAP isolates within this clade are virtually identical, and the only other sequence that is placed within the clade is HQ529498 (annotated in GenBank as “Uncultured eukaryote clone”).

### 3.2 *Poseidogoniomonas* isolates

#### 3.2.1 BEAP0338

The cells are colourless heterotrophic biflagellates of 4.6-6.1 µm long and 2.9-4.3 µm wide; two anterio-laterally oriented flagella are of a subequal length, with the anterior flagellum of 4.9-6.3 µm in length and the posterior flagellum of 4.4-5.4 µm in length (Figure 1). As a result of the flagella being roughly equal to the length of the cell and to each other, the type of movement in which the two flagella bend alternately (Suppl. Material S2) predominates over movement in which the anterior flagellum bends more actively while the posterior flagellum lies across the cell body, either passively or bending less actively compared to the anterior one.

The cells are laterally compressed and ovoid, with the two sides almost parallel to each other and slightly angularly tapered closer to the posterior end. The band of ejectosomes of 1.5-2.5 µm in length (Figure 1) was present in the majority of cells analysed. Both fully grown (just before fission) and smaller (after fission) cells often have a large, clearly visible single spherical body in the centre of the cell different from the nucleus (Figure 3-B, Figure S3).

**Figure 3.**
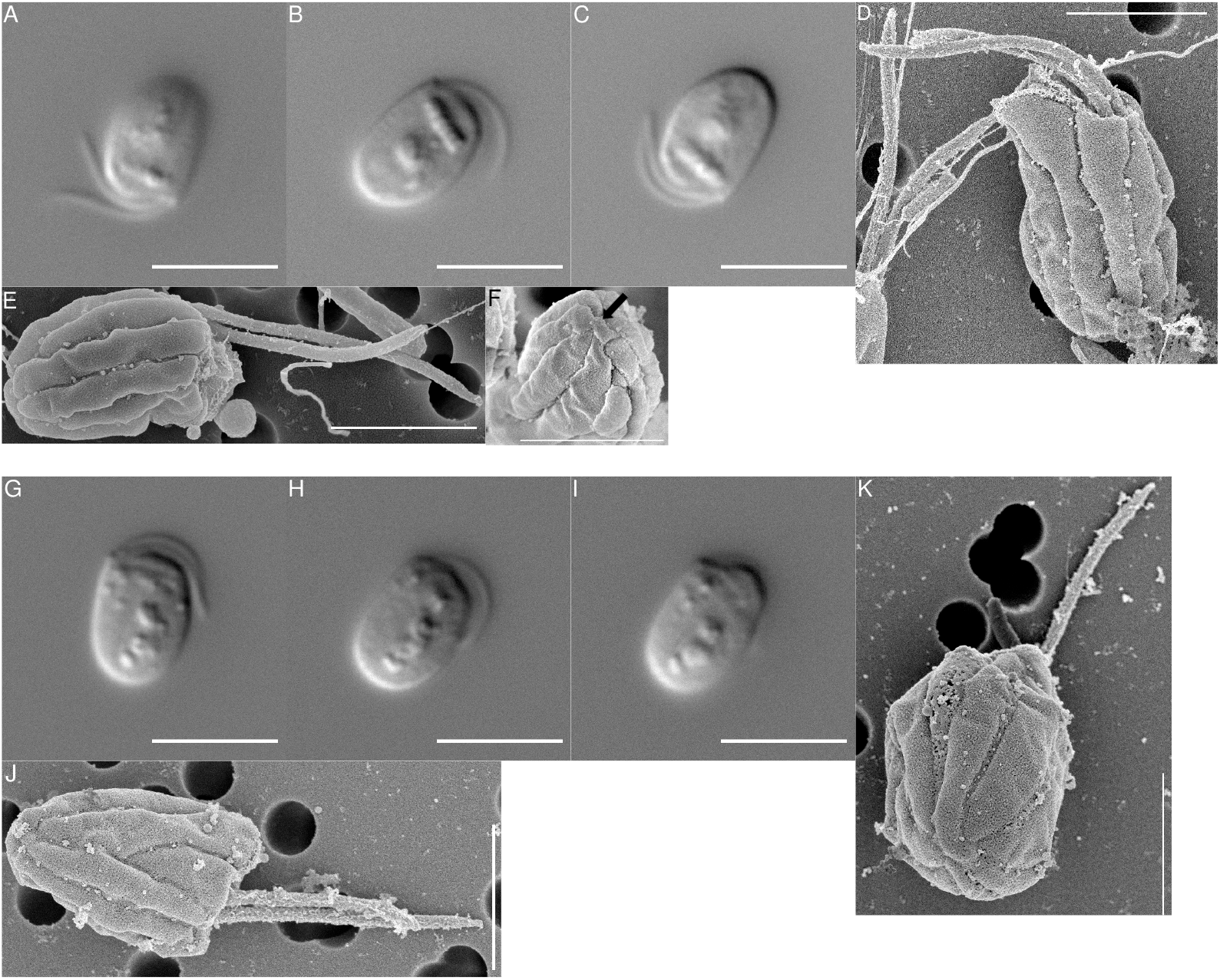
Light and electron micrographs showing *Poseidogoniomonas* isolates described in this study. A - F - *Poseidogoniomonas posteriocircus* n. sp.; G - K - *Poseidogoniomonas zopfkaesiformis* n. sp.; bar indicates 2 µm (D-F, K-J) and 5 µm (A-C, G-I); arrow in F indicates periplast plate confluence in a circular structure.

**Figure 4.**
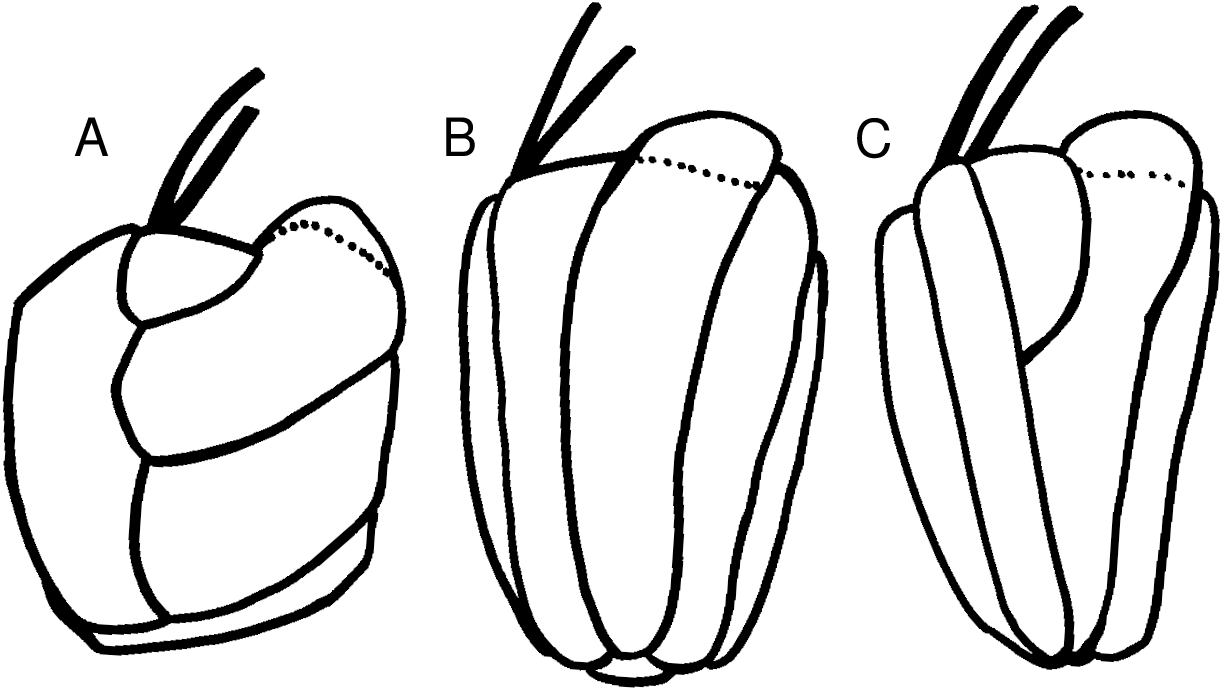
Schematic illustration of the arrangement of periplast plates for described species. A - BEAP0340 *Ebisugoniomonas hispanica* n. gen. et n. sp. (right side), B - BEAP0338 *Poseidogoniomonas posteriocircus* n. sp. (left side), C - BEAP0335 *Poseidogoniomonas zopfkaesiformis* n. sp. (left side). The dashed line shows the approximate difference of periplast plates for the other side.

In BEAP0338, five periplast plates (on the right and left sides) are arranged in parallel to one another, every plate sprawling from the anterior up to the posterior end of the cell (Figures 3-D-E, 4-B). The anterior extension which resembles what was described as a “tongue-like structure” in Phanprasert et al. (2024) is present in this strain on the left side, however it is not as prominent as in BEAP0335 (see below). The confluence of the periplast plates into a circular structure is present in this isolate (Figure 3-F); the same feature was reported for the isolate described in Martin-Cereceda et al. (2010) (*Goniomonas amphinema*, later newly combined as *Cosmogoniomonas amphinema* (Sachs et al., 2025)).

Isolates BEAP0079, BEAP0338 and BEAP0092 form a robust (99% Bootstrap, 1 posterior probability) subclade within *Poseidogoniomonas* (Figure 5). Distances between pairs of isolates within this subclade are below 0.5% (0.01 substitutions per site), while the minimal distance to the isolates from their closest sister subclade is *>*1.5% (both ML and BI) (Table 6).

**Figure 5.**
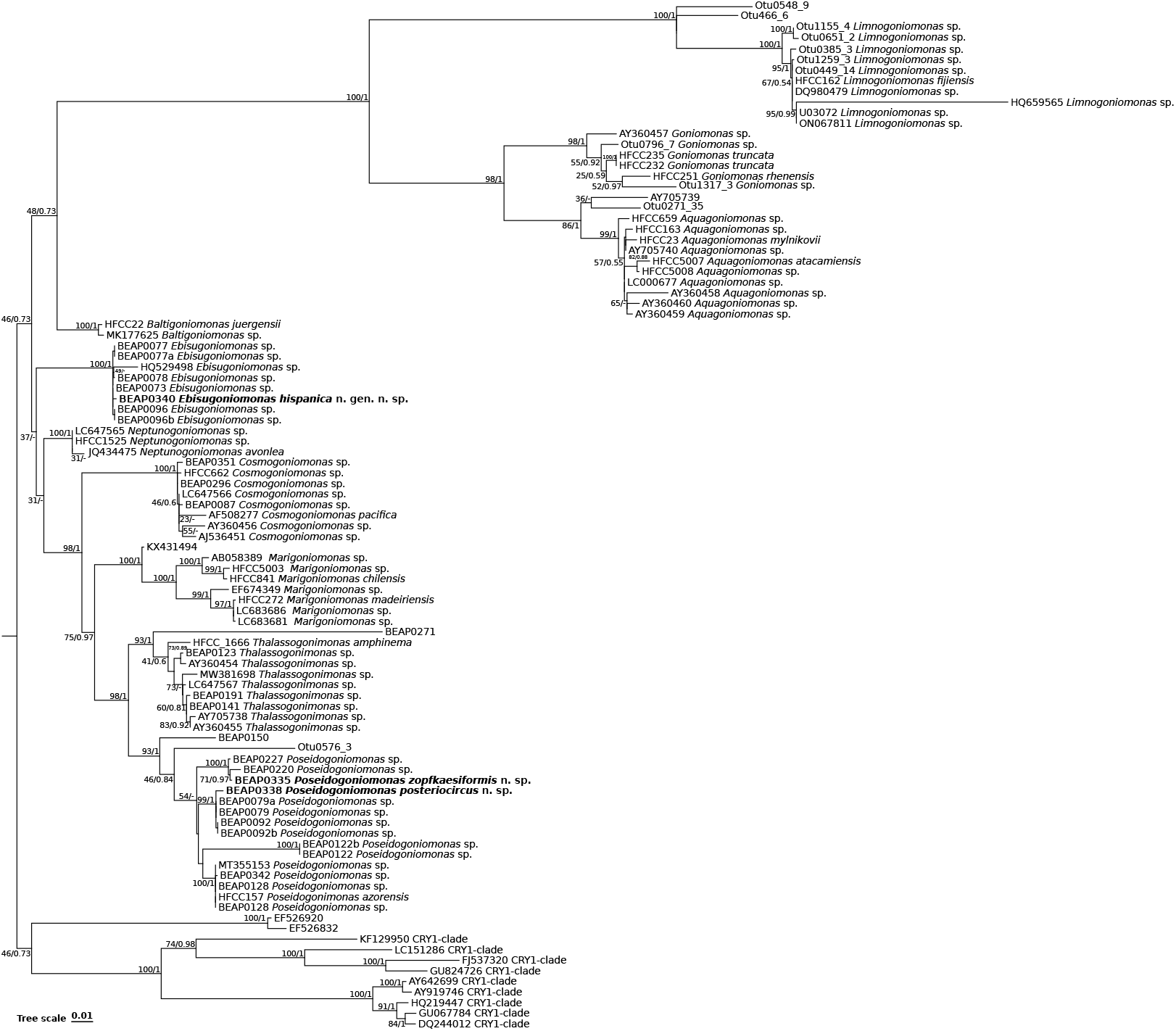
SSU rDNA Maximum Likelihood phylogeny of goniomonads. The values at the nodes represent bootstrap support and posterior probability from the Bayesian phylogeny. The values are separated by “/”, with bootstrap support first, in %, and posterior probabilities following, in decimals. For the nodes not recovered by Bayesian inference, posterior probabilities are marked as “-”. Only bootstrap supports *>*20% are shown. The scale represents substitutions per site. Bootstraps for some branches with distances *<*1% between nodes, and the *Aquagoniomonas* branch are omitted.

#### 3.2.2 BEAP0335

The cells are colourless heterotrophic biflagellates of 3.6-5.5 µm in length and 3.1-4.4 µm in width; two anterio-laterally oriented flagella are of a subunequal length, with the anterior flagellum being slightly longer (4.7-6.2 µm), and the posterior flagellum slightly shorter (4.4-5.0 µm, Figure 1). The alternate bending of both flagella is relatively frequently observed in this strain, similar to BEAP0338 (Suppl. Material S3).

The cells are laterally compressed and ovoid, with the two sides almost parallel to each other and the posterior end more rounded. The band of ejectosomes of 1-2.5 µm in length (Figure 1) was present in roughly half of all the adult cells examined.

In BEAP0335, five periplast plates (on the right and left sides) are arranged in parallel to one another. The two plates in front of the anterior depression form a conspicuous extension of the anterior part of the cell from the left side, resembling “tongue-like structure” *sensu* Phanprasert et al. (2024). *One of these two plates, which forms an extension, joins with the periplast plate that is connected with the anterior depression from which the flagella stem (Figures 3-K-J, 4-C)*. *A triangular periplast plate is present at the base of the flagellum, spanning approximately half of the cell (Figures 3-K-J, 4-C)*.

*Isolates BEAP0227, BEAP0220 and BEAP0335 form a robust (100% Bootstrap, 1 posterior probability) subclade within Poseidogoniomonas* (Figure 5). Distances between isolates within this subclade are below 1%, while the minimal distance to the isolates from the sister subclade is 2.57% (ML) or 2.41% (BI) (Table 6).

## 4 Discussion

### 4.1 Placement of the described isolates within goniomonad taxonomy

Phylogenies published in this and in previous studies (Martin-Cereceda et al., 2010; Sachs et al., 2025; Von der Heyden and Cavalier-Smith, 2004) show that the molecular diversity of goniomonads at the SSU rDNA sequence is high. For multiple lineages of protists such diverse genera were revised and split into multiple smaller genera and species based in particular on the phylogenetic affinity of investigated strains, e.g. Bendif et al. (2013); Carduck et al. (2021); Grossmann et al. (2016); Miranda Coutinho et al. (2022); Nakada et al. (2016); Schoenle et al. (2020). Given such genetic diversity, we assume that goniomonads should be split into multiple genera, and we will use the system proposed in Sachs et al. (2025) as the basis of our description.

#### 4.1.1 BEAP0340, *Ebisugoniomonas hispanica* n. gen. n. sp

The phylogenetic position of the clade with BEAP0340 is generally consistent with the one established previously in (Sachs et al., 2025, Figure 5). The clade has high support in both this and a previous (Sachs et al., 2025) study, and has >5% distance from representatives of the sister *Neptunogoniomonas* and *Baltigoniomonas* clades, which is higher than the majority of intra-genus distances observed for the representatives of marine goniomonads (*Poseidogoniomonas* being the borderline-case exception, discussed below). This indicates that *Ebisugoniomonas* n. gen. is sufficiently different from other marine/brackish goniomonad genera to be described as a new genus. The environmental sequence HQ529498 might represent a different species of the same genus.

Morphologically *Ebisugoniomonas hispanica* n. gen. n. sp. differs from the two most closely related isolates, *Baltigoniomonas* and *Neptunogoniomonas* representatives, in terms of multiple detectable features: cell length, periplast plate pattern - particularly, the angle at which plates align - and consequently cell size and shape differ from those described for both *Baltigoniomonas juergensii* and *Neptunogoniomonas avonlea* (Kim and Archibald, 2013; Sachs et al., 2025). Furthermore, the length of the posterior flagellum in relation to the cell length is different as well; the presence of toroid-shaped structures is unique.

Morphological differences from the most closely related described isolates of the sister genus, strongly supported monophyly of the clade to which *Ebisugoniomonas hispanica* n. gen. n. sp. belongs and relatively long genetic distance from the closest sister clades justifies the erection of the new genus *Ebisugoniomonas*.

#### 4.1.2 BEAP0338, *Poseidogoniomonas posteriocircus* n. sp. and BEAP0335, *Poseidogoniomonas zopfkaesiformis* n. sp

The only isolate of the *Poseidogoniomonas* genus that was previously described morphologically is HFCC157, *Poseidogoniomonas azorensis* (Sachs et al., 2025). *In this study, we collected morphological data on an isolate genetically similar (pairwise distance from HFCC157 is 0*.*08% (ML) or 0*.*19% (BI) (Table 6)) to Poseidogoniomonas azorensis* - BEAP0342. Given its morphological and genetic similarity with *Poseidogoniomonas azorensis*, we consider BEAP0342 the same species, and the data on this strain is presented in Figure S1.

In terms of morphology, strains BEAP0342, BEAP0338 and BEAP0335 differ by the length of flagella and the periplast plate pattern.

*Poseidogoniomonas posteriocircus* n. sp. has a subequal length of flagella, while *Poseidogoniomonas zopfkaesiformis* n. sp. and *Poseidogoniomonas azorensis* have posterior flagellum longer than the anterior one. The size and shape of both *Poseidogoniomonas posteriocircus* n. sp. and *Poseidogoniomonas zopfkaesiformis* n. sp. cells are similar to that of the type species of the genus *Poseidogoniomonas, Poseidogoniomonas azorensis* described in Sachs et al. (2025) and BEAP0342 (Figure S1). In *Poseidogoniomonas posteriocircus* n. sp. there is a circular structure at the posterior end of the cell (Figure 3-F), which is absent from both *Poseidogoniomonas zopfkaesiformis* n. sp. and BEAP0342 (Figure 3-K, S1-E). In both *Poseidogoniomonas azorensis* (HFCC157 and BEAP0342) and BEAP0335, a triangular periplast plate is present at the base of the flagellum, spanning approximately half of the cell (Figures 3-K-J, 4-B-C, S1-D-E). *Poseidogoniomonas posteriocircus* n. sp. lacks the triangular periplast plate (Figure 3-D-E).

Isolates BEAP0342, BEAP0338 and BEAP0335 are representatives of the three distinct well-supported subclades, which are in turn related to one another (Figure 5).

Given the ratio of pairwise distances within and between these three subclades (see Results), since previous studies (Sachs et al., 2025) generally separated representatives with a pairwise distance of >2% (which represents differences in more than 10 basepairs) into separate species, isolates BEAP0338 and BEAP0335 can be considered as new species within *Poseidogoniomonas*. The erection of new species is further supported by the morphological differences distinguishing *Poseidogoniomonas posteriocircus* n. sp. and *Poseidogoniomonas zopfkaesiformis* n. sp. from each other and from *Poseidogoniomonas azorensis* with BEAP0342.

Finally, the assemblage of monophyletic lineages that are consistently recovered as an ingroup relative to strain BEAP0150 has high genetic diversity within itself, with four well-supported monophyletic clades that have ambiguous branching order among them. Here we describe the representatives from two of these lineages as *Poseidogoniomonas* species, assuming that *Poseidogoniomonas* is a monophyletic clade that is sister the sequence of the isolate BEAP0150. We do this in particular to avoid possible confusion in the future, as adding more sequences to this clade might result in the resolution of what currently resembles a polyphyletic relationship between the four stable clades. If each of the four clades is described as a novel genus, there is a probability such genera would have to be revisited in the future.

### 4.2 General considerations on morphology

All the species now described in the genus *Poseidogoniomonas* have virtually identical cell sizes and shapes, but the length of flagella and the periplast plate arrangement pattern can be used to distinguish the previously described *Poseidogoniomonas azorensis* from *Poseidogoniomonas posteriocircus* n. sp. and *Poseidogoniomonas zopfkaesiformis* n. sp.

The variation in the number of periplast plates is limited: in the isolates investigated in this study, as well as in goniomonad isolates described previously, it consistently fluctuates between four and six (here we define every periplast striation as a “plate”). In contrast, the pattern in which periplast plates are arranged has a high potential to serve as a taxonomically important character which can be used for discriminating species from one another. This pattern is conserved across multiple cells of the same strain (Phanprasert et al., 2024) and does not appear to change depending on culture growth stage or sample preparation (verified in this study, data not shown). The presence of the triangular periplast plates at the flagellar base, the angle at which the plates are oriented towards one another and presence of the periplast plate confluence in a circular structure are particularly interesting features for morphological characterisation.

We also note three additional features: (1) the toroid structures at the anterior end of the cell in *Ebisugoniomonas hispanica* n. gen. n. sp.; (2) the differing predominant movement patterns among the isolates described, which are most likely explained by flagella lengths (a similar observation was reported in Kim and Archibald, 2013); (3) a varying frequency of ejectosome row presence/absence in different strains.

All the aforementioned accessory features observed in the studied strains and the fact that these features are strain-specific provoke important questions on the biology and ecology of goniomonads. Can different movement patterns affect the feeding strategies and/or habitat preferences of goniomonads? Does the frequency of presence/absence of ejectosome rows in cells depend on environmental factors (previously addressed in Schuster (1968) but not subsequently investigated), ejectosome turnover rates, the age/density of the population, or other factors? Addressing those questions becomes even more interesting when considering that all the strains investigated here were isolated from the same location. The possibility that they coexist in the same habitat leads us to speculate that the differences in their cell biology, morphology and behaviours may have been the causes for their ecological niche separation and may have allowed sympatric speciation.

Overall, our knowledge of cell biology, behaviour and morphology of this group remains extremely limited, yet detailed studies on multiple goniomonad species can provide valuable insights into how these species live and function, how they differ from one another, and possibly, how they specialise and adapt to specific environmental factors that are different among closely related species.

### 4.3 General considerations on the phylogeny of goniomonads

Compared to the recent study on goniomonad phylogeny published in Sachs et al. (2025), this study included additional goniomonad sequences from three sources: (1) environmental sequences from long-read SSU rDNA sequencing dataset from Jamy et al. (2020), (2) a few environmental sequences from NCBI nt database which were not classified as goniomonads but were instead retrieved by an iterative search versus NCBI nt using a part of EukRef pipeline and (3) the sequences of goniomonad isolates that we obtained after the phylogeny given by Sachs et al. (2025) was released.

With an expanded dataset of partial SSU rDNA sequences, several changes in the topology of the goniomonad phylogenetic tree were observed when compared to that of Sachs et al. (2025); some of those are not directly related to the genera investigated here.

In the freshwater goniomonad clade, environmental sequences from the long-read dataset of Jamy et al. (2020); OTU0796_7 and OTU1317_3 indicate that there are more related species within the genus *Goniomonas*, while OTU1155_4 and OTU0651_2 indicate a higher diversity in the *Limnogoniomonas* clade. More remarkable are the positions of the environmental sequences OTU0548_9 and OTU466_6 which are sister to the *Limnogoniomonas* clade. Similarly, OTU0271_35 is sister to the clade representing the *Aquagoniomonas* genus.

The environmental sequence OTU0576_3 that was initially recovered as a part of long-read dataset (Jamy et al., 2020) (according to the metadata, this sequence belongs to a marine sample), clustered within the marine genus of *Poseidogoniomonas*, with low (46%) support according to ML but high (0.84) posterior probability according to BI. Manual inspection of the alignment did not reveal any evident artefacts in this sequence (i.e., long unaligned regions), which is why this sequence was retained.

For marine goniomonads, there is a minor discrepancy in the branching order of *Ebisugoniomonas* n. gen. in the phylogenies of the current study compared to the one published previously in Sachs et al. (2025). In our phylogeny, the *Ebisugoniomonas* clade branches between (i) *Baltigoniomonas* sequences MK177625 and HFCC22 and (ii) *Neptunogoniomonas* sequences JQ434475, HFCC1525 and LC647565.

In addition, the BEAP0150 sequence (from a culture which was lost before morphological studies could be conducted) branches as a sister lineage to all *Poseidogoniomonas* sequences. Another enigmatic long branch, this time sister to the *Thalassogoniomonas* clade belongs to BEAP0271 (another culture that was lost prior to morphological studies). Finally, the environmental sequences EF674349 and AB058389, as well as AJ536451 and AY360456 indicate the presence of further undescribed species of *Marigoniomonas* and *Cosmogoniomonas*, respectively (based on their pairwise distances to other representatives of the genus).

Environmental sequences EF526920 and EF526832 are sister to the CRY-1 environmental lineage, and, on the other side, to all the goniomonads. Given the low support for the node in which EF526920 and EF526832 diverge from CRY-1 (42% bootstrap and 0.61 posterior probability), these two sequences might as well represent a very basal lineage of goniomonads (Figures 5 and S4). The position of this clade also remains stable when more sequences are added as an outgroup (Figures S5 and S6).

Overall, the phylogenetic position of the previously unknown or undetected goniomonad sequences which were added to the dataset from Sachs et al. (2025) suggests that the molecular diversity of goniomonads still remains understudied. The more sequences are obtained, both from environmental DNA and cultured isolates, the higher is the chance to find early branching or novel clades. This highlights the need for further exploration of the astonishing diversity of goniomonads.

### 4.4 Concluding remarks and outlook

*Ebisugoniomonas hispanica* n. gen. n. sp. is a representative species of a novel goniomonad genus that has an early-branching position within the goniomonad phylogeny. *Poseidogoniomonas posteriocircus* n. sp. and *Poseidogoniomonas zopfkaesiformis* n. sp. are two novel species of the recently described genus *Poseidogoniomonas*. They differ both genetically and morphologically from a third *Poseidogoniomonas* representative, *Poseidogoniomonas azorensis*. The relative arrangement of periplast plates appears to be a potentially important morphological feature complex that can be used for subsequent discrimination of species based on morphology. Light microscopy methods, applied to goniomonads described here, revealed additional details on the range of cellular structures: we attempted to establish the nature of globules in the cytoplasm of *Ebisugoniomonas hispanica* n. gen. n. sp. and *Poseidogoniomonas posteriocircus* n. sp. Goniomonad cells of different species can be distinguished morphologically, and comparative studies of goniomonad morphology could serve as an important basis for future functional studies of various species within this lineage. The phylogeny of goniomonads, expanded by the addition of novel sequences, reveals the existence of multiple genus- and species-level lineages within this clade and indicates that our knowledge of goniomonad genetic diversity is far from being exhaustive and is likely to expand as more data becomes available.

## 5 TAXONOMIC SUMMARY

Assignment: Eukarya, Cryptista, Cryptophyta, Goniomonadea, Goniomonadida

### *Ebisugoniomonas* Zavadska and Richter, n. gen

#### Diagnosis

Biflagellate, colourless, unicellular cryptist. Cells laterally flattened and round. The anterior flagellum is shorter than the posterior one. The posterior flagellum is most commonly longer than the cell body. Five periplast plates on both sides.

#### Type species

*Ebisugoniomonas hispanica* n.sp.

#### Etymology

Ebisu is a Japanese deity of fishery and luck, associated with marine fauna. *Ebisugoniomonas* continues the practice of marine genera of gonimonads named after sea deities (e.g. *Neptunogoniomonas, Poseidogoniomonas*).

#### Type species

*Ebisugoniomonas hispanica* n. gen. et. n. sp.

### *Ebisugoniomonas hispanica* Zavadska and Richter, n. sp

#### Diagnosis

Free-living, colourless, marine unicellular biflagellates with laterally compressed cells. Cells are close to round, with length (4.8-6.4 µm, mean=5.5 µm, sd=0.37, n=30) nearly equal to the width (4.3-5.6 µm, mean=4.8 µm, sd=0.38, n=30). The two flagella arising from an anterior depression are unequal in length. The anterior flagellum is 5.5-7.5 µm long (mean = 6.3 µm, sd = 0.4, n=25) and the posterior flagellum is 8.3-10.2µm long (mean=9.0 µm, sd=0.51, n=13). A band of ejectosomes is occasionally present at the anterior side of the cell, 1.3-2.9µm in length (mean=2.2 µm, sd=0.43, n=13). Cells carry five periplast plates on the right and left sides; the triangular-shaped plate is located at the base of the flagella.

#### Type location

Strain was obtained from seawater at a depth of 1 meter in the Mediterranean Sea near Blanes Bay, Spain, 41°40’N 2°40’E.

#### Type material

The culture of the type strain is deposited in the Roscoff Culture Collection, Roscoff Biological Station, National Center of Scientific Research, Sorbonne University, Roscoff, under ID 11327. The SEM stub is deposited in the Marine Biological Reference Collection of the Institute of Marine Science, Spanish Academy of Sciences, Barcelona, under ID XXX (will be provided prior to publication) (Santos-Bethencourt et al., 2023).

#### Gene sequence data

SSU rDNA sequence from the isolate BEAP0340 was deposited in Genbank under ID PV090905.

#### ZooBank ID

XXX (will be provided prior to publication)

#### Etymology

“Hispanica” refers to the site of isolation which is located in Spain.

### *Poseidogoniomonas posteriocircus* Zavadska and Richter, n. sp

#### Diagnosis

Free-living, colourless, marine unicellular biflagellates with laterally compressed cells. Cells are ovoid, with two longer sides almost parallel to each other, tapered at the posterior end. Cell length (4.6-6.1 µm, mean=5.2 µm, sd=0.42, n=33) is larger than the width (2.9-4.3 µm, mean=3.7 µm, sd=0.32, n=33). The two flagella arising from an anterior depression are subequal in length. The anterior flagellum is 4.9-6.3 µm long (mean = 5.5 µm, sd = 0.3, n=21) and the posterior flagellum is 4.4-5.4µm long (mean=4.8 µm, sd=0.3, n=13). A band of ejectosomes can be present at the anterior side of the cell, 1.6-2.5µm in length (mean=2.2 µm, sd=0.23, n=23). Cells carry five periplast plates on the right and left sides; the triangular-shaped plate at the base of the flagella is absent; the periplast plates converge into a circular structure at the posterior end of the cell.

#### Type location

Strain was obtained from seawater at a depth of 1 meter in the Mediterranean Sea near Blanes Bay, Spain, 41°40’N 2°40’E.

#### Type material

The culture of the type strain is deposited in the Roscoff Culture Collection, Roscoff Biological Station, National Center of Scientific Research, Sorbonne University, Roscoff, under ID 11329. The SEM stub is deposited in the Marine Biological Reference Collection of the Institute of Marine Science, Spanish Academy of Sciences, Barcelona, under ID XXX (will be provided prior to publication) (Santos-Bethencourt et al., 2023).

#### Gene sequence data

SSU rDNA sequence from the isolate BEAP0338 was deposited in Genbank under ID PV090907.

#### ZooBank ID

XXX (will be provided prior to publication)

#### Etymology

“Posteriocircus” refers to the circular structure present at the posterior end of the cell.

### *Poseidogoniomonas zopfkaesiformis* Zavadska and Richter, n. sp

#### Diagnosis

Free-living, colourless, marine unicellular biflagellates with laterally compressed cells. Cells are ovoid, with two longer sides almost parallel to each other, rounded at the posterior end. Cell length (4.0-5.5 µm, mean=4.8 µm, sd=0.42, n=32) is larger than the width (3.3-4.4 µm, mean=3.7 µm, sd=0.3, n=32). The two flagella arising from an anterior depression are slightly different in length. The anterior flagellum (4.7-6.2 µm, mean = 5.5 µm, sd = 0.36, n=24) is slightly longer than the posterior flagellum (4.4-5.0µm, mean=4.7 µm, sd=0.2, n=18). A band of ejectosomes is occasionally present at the anterior side of the cell, 1.2-2.5µm in length (mean=1.7 µm, sd=0.43, n=6). Cells carry five periplast plates on right and left sides; the triangular-shaped plate is present at the base of the flagella.

#### Type location

Strain was obtained from seawater at a depth of 1 meter in the Mediterranean Sea near Blanes Bay, Spain, 41°40’N 2°40’E.

#### Type material

The culture of the type strain is deposited in the Roscoff Culture Collection, Roscoff Biological Station, National Center of Scientific Research, Sorbonne University, Roscoff, under ID 11333. The SEM stub is deposited in the Marine Biological Reference Collection of the Institute of Marine Science, Spanish Academy of Sciences, Barcelona, under ID XXX (will be provided prior to publication) (Santos-Bethencourt et al., 2023).

#### Gene sequence data

SSU rDNA sequence from the isolate BEAP0335 was deposited in Genbank under ID PV090904.

#### ZooBank ID

XXX (will be provided prior to publication)

#### Etymology

“Zopfkäse” is a type of cheese which visually resembles the periplast plate pattern observed in goniomonads, and in particular in this species.

## 6 Acknowledgements

We thank Maria Carlos Oliveira for keeping the mixed culture with *Ebisugoniomonas* alive. We also thank other members of BEAP lab - Àlex Gàlvez i Morante and Margarita Skamnelou for kindly providing cultures and sequences of goniomonads, Xènia Maya for maintaining the described isolates. We thank Cédric Berney for discussions on SSU rDNA alignments. We thank to Rosita Bieg, Brigitte Gräfe and Anke Pyschny for their advice on the technical details of protocols applied to goniomonads. We thank Electron Microscopy facility, and heads of the facility, Mariona Segura and José Manuel Fortuño for all the technical support with SEM imaging, reliability and advice on sample preparation. This work was supported by the grant QC2021-007134-P funded by MCIN/AEI/10.13039/501100011033 and by the “European Union NextGenerationEU/PRTR” for electron microscopy, the European Research Council (ERC) under the European Union’s Horizon 2020 research and innovation programme (grant agreement No. 949745), by grant PID2023-152955NA-I00 funded by MICIU/AEI/10.13039/501100011033 and by ERDF/EU and the Departament de Recerca i Universitats de la Generalitat de Catalunya (exp. 2021 SGR 00751).G H I K

## References

S. M. Adl, D. Bass, C. E. Lane, J. Lukeš, C. L. Schoch, A. Smirnov, S. Agatha, C. Berney, M. W. Brown, F. Burki, et al. Revisions to the classification, nomenclature, and diversity of eukaryotes. Journal of Eukaryotic Microbiology, 66(1):4–119, 2019.

E. M. Bendif, I. Probert, D. C. Schroeder, and C. de Vargas. On the description of Tisochrysis lutea gen. nov. sp. nov. and Isochrysis nuda sp. nov. in the Isochrysidales, and the transfer of Dicrateria to the Prymnesiales (Haptophyta). Journal of applied phycology, 25:1763–1776, 2013.

C. Berney, N. Henry, F. Mahé, D. J. Richter, and C. de Vargas. EukRibo: a manually curated eukaryotic 18S rDNA reference database to facilitate identification of new diversity. bioRxiv, 2022. doi: 10.1101/2022.11.03.515105. URL https://www.biorxiv.org/content/early/2022/11/04/2022.11.03.515105.

S. Capella-Gutiérrez, J.M. Silla-Martínez, and T. Gabaldón. trimAl: a tool for automated alignment trimming in large-scale phylogenetic analyses. Bioinformatics, 25(15):1972–1973, 06 2009. ISSN 1367-4803. doi: 10.1093/bioinformatics/btp348. URL https://doi.org/10.1093/bioinformatics/btp348.

Carduck, F. Nitsche, A. Rybarski, M. Hohlfeld, and H. Arndt. Diversity and phylogeny of percolomonads based on newly discovered species from hypersaline and marine waters. European Journal of Protistology, 80:125808, 2021.

Cavalier-Smith. Principles of Protein and Lipid Targeting in Secondary Symbiogenesis: Euglenoid, Dinoflagellate, and Sporozoan Plastid Origins and the Eukaryote Family Tree. Journal of Eukaryotic Microbiology, 46(4):347–366, 1999. doi: 10.1111/j.1550-7408.1999.tb04614.x. URL https://onlinelibrary.wiley.com/doi/abs/10.1111/j.1550-7408.1999.tb04614.x.

Cenci, S. J. Sibbald, B. A. Curtis, R. Kamikawa, L. Eme, D. Moog, B. Henrissat, E. Maréchal, M. Chabi, C. Djemiel, et al. Nuclear genome sequence of the plastid-lacking cryptomonad Goniomonas avonlea provides insights into the evolution of secondary plastids. BMC biology, 16:1–23, 2018.

B. L. Clay. Chapter 18 - cryptomonads. In J. D. Wehr, R. G. Sheath, and J. P. Kociolek, editors, Freshwater Algae of North America (Second Edition), Aquatic Ecology, pages 809–850. Academic Press, Boston, second edition edition, 2015. ISBN 978-0-12-385876-4. doi: 10.1016/B978-0-12-385876-4.00018-9. URL https://www.sciencedirect.com/science/article/pii/B9780123858764000189.

D. Darriba, D. Posada, A. M. Kozlov, A. Stamatakis, B. Morel, and T. Flouri. ModelTest-NG: A New and Scalable Tool for the Selection of DNA and Protein Evolutionary Models. Molecular Biology and Evolution, 37(1):291–294, 08 2019. ISSN 0737-4038. doi: 10.1093/molbev/msz189. URL https://doi.org/10.1093/molbev/msz189.

J. A. Deane, I. M. Strachan, G. W. Saunders, D. R. A. Hill, and G. I. McFadden. Cryptomonad evolution: nuclear 18S rDNA phylogeny versus cell morphology and pigmentation. Journal of Phycology, 38(6):1236–1244, 2002. doi: 10.1046/j.1529-8817.2002.01250.x. URL https://onlinelibrary.wiley.com/doi/abs/10.1046/j.1529-8817.2002.01250.x.

J. del Campo, M. Kolisko, V. Boscaro, L. F. Santoferrara, S. Nenarokov, R. Massana, L. Guillou, A. Simpson, C. Berney, C. de Vargas, M. W. Brown, P. J. Keeling, and L. Wegener Parfrey. EukRef: Phylogenetic curation of ribosomal RNA to enhance understanding of eukaryotic diversity and distribution. PLOS Biology, 16(9):1–14, 09 2018. doi: 10.1371/journal.pbio.2005849. URL https://doi.org/10.1371/journal.pbio.2005849.

O. Flegontova, P. Flegontov, S. Malviya, J. Poulain, C. de Vargas, C. Bowler, J. Lukeš, and A. Horák. Neobodonids are dominant kinetoplastids in the global ocean. Environmental Microbiology, 20(2):878–889, 2018. doi: 10.1111/1462-2920.14034. URL https://enviromicro-journals.onlinelibrary.wiley.com/doi/abs/10.1111/1462-2920.14034.

L. Grossmann, C. Bock, M. Schweikert, and J. Boenigk. Small but Manifold – Hidden Diversity in “Spumella-like Flagellates”. Journal of Eukaryotic Microbiology, 63(4):419–439, 2016. doi: 10.1111/jeu.12287. URL https://onlinelibrary.wiley.com/doi/abs/10.1111/jeu.12287.

K. Hoef-Emden and J. M. Archibald. Cryptophyta (Cryptomonads), pages 851–891. Springer International Publishing, Cham, 2017. ISBN 978-3-319-28149-0. doi: 10.1007/978-3-319-28149-0_35. URL https://doi.org/10.1007/978-3-319-28149-0_35.

M. Jamy, R. Foster, P. Barbera, L. Czech, A. Kozlov, A. Stamatakis, G. Bending, S. Hilton, D. Bass, and F. Burki. Long-read metabarcoding of the eukaryotic rDNA operon to phylogenetically and taxonomically resolve environmental diversity. Molecular Ecology Resources, 20(2):429–443, 2020. doi: 10.1111/1755-0998.13117. URL https://onlinelibrary.wiley.com/doi/abs/10.1111/1755-0998.13117.

K. Katoh, K. Misawa, K. Kuma, and T. Miyata. MAFFT: a novel method for rapid multiple sequence alignment based on fast Fourier transform. Nucleic Acids Research, 30(14):3059–3066, 07 2002. ISSN 0305-1048. doi: 10.1093/nar/gkf436. URL https://doi.org/10.1093/nar/gkf436.

E. Kim and J. M. Archibald. Ultrastructure and Molecular Phylogeny of the Cryptomonad Goniomonas avonlea sp. nov. Protist, 164(2):160–182, 2013. ISSN 1434-4610. doi: 10.1016/j.protis.2012.10.002. URL https://www.sciencedirect.com/science/article/pii/S1434461012001009.

J. Larsen and D. J. Patterson. Some flagellates (Protista) from tropical marine sediments. Journal of Natural History, 24 (4):801–937, 1990. doi: 10.1080/00222939000770571. URL https://doi.org/10.1080/00222939000770571.

P. Lopez-Garcia, H. Philippe, F. Gail, and D. Moreira. Autochthonous eukaryotic diversity in hydrothermal sediment and experimental microcolonizers at the Mid-Atlantic Ridge. Proceedings of the National Academy of Sciences of the United States of America, 100:697–702, 02 2003. doi: 10.1073/pnas.0235779100.

F. Madeira, N. Madhusoodanan, J. Lee, A. Eusebi, A. Niewielska, A. R. N. Tivey, R. Lopez, and S. Butcher. The EMBL-EBI Job Dispatcher sequence analysis tools framework in 2024. Nucleic acids research, 52(W1):W521—W525, July 2024. ISSN 0305-1048. doi: 10.1093/nar/gkae241. URL https://europepmc.org/articles/PMC11223882.

B. Marin, M. Klingberg, and M. Melkonian. Phylogenetic Relationships among the Cryptophyta: Analyses of Nuclear-Encoded SSU rRNA Sequences Support the Monophyly of Extant Plastid-Containing Lineages. Protist, 149(3):265–276, 1998. ISSN 1434-4610. doi: 10.1016/S1434-4610(98)70033-1. URL https://www.sciencedirect.com/science/article/pii/S1434461098700331.

M. Martin-Cereceda, E. C. Roberts, E. C. Wootton, E. Bonaccorso, P. Dyal, A. Guinea, D. Rogers, C. J. Wright, and G. Novarino. Morphology, ultrastructure, and small subunit rDNA phylogeny of the marine heterotrophic flagellate Goniomonas aff. amphinema. Journal of Eukaryotic Microbiology, 57(2):159–170, 2010.

G. I. McFadden. Genome of tiny predator with big appetite. BMC biology, 16(1):140, 2018.

G. P. McFadden, G.I. and D. Hill. Goniomonas: rRNA sequences indicate that this phagotrophic flagellate is a close relative of the host component of cryptomonads. European Journal of Phycology, 29(1):29–32, 1994. doi: 10.1080/09670269400650451. URL https://doi.org/10.1080/09670269400650451.

J. P. Mignot. Étude ultrastructurale de (Cyathomonas truncata) From.(flagellé cryptomonadine). 1965.

L. Miranda Coutinho, F. Penelas Gomes, M. Nasri Sissini, T. Vieira-Pinto, M. C. Muller de Oliveira Henriques, M. C. Oliveira, P. Antunes Horta, and M. B. Barbosa de Barros Barreto. Cryptic diversity in non-geniculate coralline algae: a new genus Roseolithon (Hapalidiales, Rhodophyta) and seven new species from the Western Atlantic. European Journal of Phycology, 57(2):227–250, 2022.

T. Nakada, M. Tomita, J.-T. Wu, and H. Nozaki. Taxonomic revision of Chlamydomonas subg. Amphichloris (Volvocales, Chlorophyceae), with resurrection of the genus Dangeardinia and descriptions of Ixipapillifera gen. nov. and Rhysamphichloris gen. nov. Journal of Phycology, 52(2):283–304, 2016. doi: 10.1111/jpy.12397. URL https://onlinelibrary.wiley.com/doi/abs/10.1111/jpy.12397.

G. Novarino. A companion to the identification of cryptomonad flagellates (Cryptophyceae= Cryptomonadea). In Phytoplankton and Equilibrium Concept: The Ecology of Steady-State Assemblages: Proceedings of the 13th Workshop of the International Association of Phytoplankton Taxonomy and Ecology (IAP), held in Castelbuono, Italy, 1–8 September 2002, pages 225–270. Springer, 2003.

J. A. Packer, D. Zavadska, E. J. Weston, Y. Eglit, D. J. Richter, and A. G. B. Simpson. Characterization of Allobodo yubaba sp. nov. and Novijibodo darinka gen. et sp. nov., cultivable free-living species of the phylogenetically enigmatic kinetoplastid taxon Allobodonidae. Journal of Eukaryotic Microbiology, 72(1):e13072, 2025. doi: 10.1111/jeu.13072. URL https://onlinelibrary.wiley.com/doi/abs/10.1111/jeu.13072.e13072JEUKMIC-24-5791.

D. J. Patterson and J. Larsen. The Biology of Free-Living Heterotrophic Flagellates. Oxford University Press, 01 1992. ISBN 9780198577478. doi: 10.1093/oso/9780198577478.001.0001. URL https://doi.org/10.1093/oso/9780198577478.001.0001.

Y. Phanprasert, S. Y. Kim, N. S. Kang, M. Jeong, J. I. Kim, W. Shin, W. J. Lee, and E. Kim. Morphological and Molecular Phylogenetic Characterization of Three New Marine Goniomonad Species. bioRxiv, 2024. doi: 10.1101/2024.12.12.627435. URL https://www.biorxiv.org/content/early/2024/12/13/2024.12.12.627435.

K. Piwosz, J. Kownacka, A. Ameryk, M. Zalewski, and J. Pernthaler. Phenology of cryptomonads and the CRY1 lineage in a coastal brackish lagoon (Vistula Lagoon, Baltic Sea). Journal of Phycology, 52(4):626–637, 2016. doi: 10.1111/jpy.12424. URL https://onlinelibrary.wiley.com/doi/abs/10.1111/jpy.12424.

M. N. Price, P. S. Dehal, and A. P. Arkin. FastTree 2 – Approximately Maximum-Likelihood Trees for Large Alignments. PLOS ONE, 5(3):1–10, 03 2010. doi: 10.1371/journal.pone.0009490. URL https://doi.org/10.1371/journal.pone.0009490.

D. J. Richter, P. Fozouni, M. B. Eisen, and N. King. Gene family innovation, conservation and loss on the animal stem lineage. eLife, 7:e34226, may 2018. ISSN 2050-084X. doi: 10.7554/eLife.34226. URL https://doi.org/10.7554/eLife.34226.

F. Ronquist and J. P. Huelsenbeck. MrBayes 3: Bayesian phylogenetic inference under mixed models. Bioinformatics, 19(12):1572–1574, 08 2003. ISSN 1367-4803. doi: 10.1093/bioinformatics/btg180. URL https://doi.org/10.1093/bioinformatics/btg180.

M. Sachs, F. Nitsche, and H. Arndt. Cryptic Cryptophytes-revision of the genus Goniomonas. bioRxiv, 2025. doi: 10.1101/2024.07.17.603845. URL https://www.biorxiv.org/content/early/2025/02/05/2024.07.17.603845.

R. Santos-Bethencourt, A. Sabatés, M. Ramón, R. Villanueva, A. Lombarte, P. Abelló, and E. Guerrero. Marine Biological Reference Collections: CBMR-General (ICM-CSIC). 2023.

J. Schindelin, I. Arganda-Carreras, E. Frise, V. Kaynig, M. Longair, T. Pietzsch, S. Preibisch, C. Rueden, S. Saalfeld,B. Schmid, J.-Y. Tinevez, D. J. White, V. Hartenstein, K. Eliceiri, P. Tomancak, and A. Cardona. Fiji: an open-source platform for biological-image analysis. Nature Methods, 9(7):676–682, Jul 2012. ISSN 1548-7105. doi: 10.1038/nmeth.2019. URL https://doi.org/10.1038/nmeth.2019.

A. Schoenle, M. Hohlfeld, M. Rosse, P. Filz, C. Wylezich, F. Nitsche, and H. Arndt. Global comparison of bicosoecid Cafeteria-like flagellates from the deep ocean and surface waters, with reorganization of the family Cafeteriaceae. European Journal of Protistology, 73:125665, 2020. ISSN 0932-4739. doi: 10.1016/j.ejop.2019.125665. URL https://www.sciencedirect.com/science/article/pii/S0932473919301026.

F. Schuster. The gullet and trichocysts of Cyathomonas truncata. Experimental Cell Research, 49(2):277–284, 1968. ISSN 0014-4827. doi: 10.1016/0014-4827(68)90178-X. URL https://www.sciencedirect.com/science/article/pii/001448276890178X.

K. Shalchian-Tabrizi, J. Bråte, R. Logares, D. Klaveness, C. Berney, and K. S. Jakobsen. Diversification of unicellular eukaryotes: cryptomonad colonizations of marine and fresh waters inferred from revised 18S rRNA phylogeny. Environmental Microbiology, 10(10):2635–2644, 2008. doi: 10.1111/j.1462-2920.2008.01685.x. URL https://enviromicro-journals.onlinelibrary.wiley.com/doi/abs/10.1111/j.1462-2920.2008.01685.x.

W. Shen, S. Le, Y. Li, and F. Hu. SeqKit: A Cross-Platform and Ultrafast Toolkit for FASTA/Q File Manipulation. PLOS ONE, 11(10):1–10, 10 2016. doi: 10.1371/journal.pone.0163962. URL https://doi.org/10.1371/journal.pone.0163962.

J. L. Steenwyk, I. Buida, Thomas J, A. L. Labella, Y. Li, X.-X. Shen, and A. Rokas. PhyKIT: a broadly applicable UNIX shell toolkit for processing and analyzing phylogenomic data. Bioinformatics, 37(16):2325–2331, 02 2021. ISSN 1367-4803. doi: 10.1093/bioinformatics/btab096. URL https://doi.org/10.1093/bioinformatics/btab096.

G. Tanifuji and N. T. Onodera. Chapter Eight - Cryptomonads: A Model Organism Sheds Light on the Evolutionary History of Genome Reorganization in Secondary Endosymbioses. In Y. Hirakawa, editor, Secondary Endosymbioses, volume 84 of Advances in Botanical Research, pages 263–320. Academic Press, 2017. doi: 10.1016/bs.abr.2017.06.005. URL https://www.sciencedirect.com/science/article/pii/S0065229617300526.

A. Vardanian. USearch by Unum Cloud, Oct. 2023. URL https://github.com/unum-cloud/usearch.

D. Vaulot, S. Geisen, F. Mahé, and D. Bass. PR2-primers: An 18S rRNA primer database for protists. Molecular Ecology Resources, 22(1):168–179, 2022. doi: 10.1111/1755-0998.13465. URL https://onlinelibrary.wiley.com/doi/abs/10.1111/1755-0998.13465.

E. C. Von der Heyden, S. and T. Cavalier-Smith. Genetic diversity of goniomonads: an ancient divergence between marine and freshwater species. European Journal of Phycology, 39(4):343–350, 2004. doi: 10.1080/09670260400005567. URL https://doi.org/10.1080/09670260400005567.

A. M. Waterhouse, J. B. Procter, D. M. A. Martin, M. Clamp, and G. J. Barton. Jalview Version 2—a multiple sequence alignment editor and analysis workbench. Bioinformatics, 25(9):1189–1191, 01 2009. ISSN 1367-4803. doi: 10.1093/bioinformatics/btp033. URL https://doi.org/10.1093/bioinformatics/btp033.

J. Xie, Y. Chen, G. Cai, R. Cai, Z. Hu, and H. Wang. Tree Visualization By One Table (tvBOT): a web application for visualizing, modifying and annotating phylogenetic trees. Nucleic Acids Research, 51(W1):W587–W592, 05 2023. ISSN 0305-1048. doi: 10.1093/nar/gkad359. URL https://doi.org/10.1093/nar/gkad359.

A. Yabuki, R. Kamikawa, S. A. Ishikawa, M. Kolisko, E. Kim, A. S. Tanabe, K. Kume, K.-i. Ishida, and Y. Inagki. Palpitomonas bilix represents a basal cryptist lineage: insight into the character evolution in Cryptista. Scientific reports, 4(1):4641, 2014.

