## Supplemental information for "The morphology and small subunit rDNA gene phylogeny of the novel goniomonad genus *Ebisugoniomonas* and two novel *Poseidogoniomonas* species"

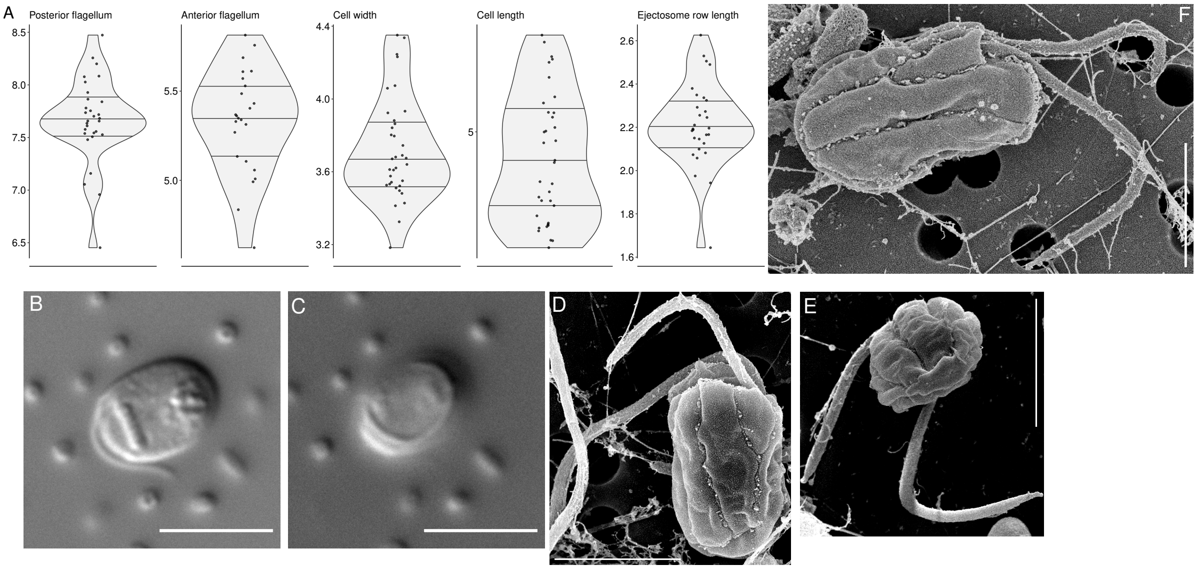


Figure S1: **BEAP0342, the strain genetically close to *Poseidogoniomonas azorensis*.** A - Summary of morphological parameters; B-F - Light and electron micrographs showing BEAP0342 isolate; bar indicates 2 µm (D-E) and 5 µm (B-C); Note the absence of the circular structure at the periplast plate confluence in E. BEAP0342 is the monoclonal culture derived from the strain BEAP0128. BEAP0342 has a cophenetic distance of 0.08% (ML) or 0.19% (BI) from strain HFCC157 (HFCC157 strain is described as *P. azorensis* in (Sachs et al., 2025)), and is similar to *P. azorensis* by morphological parameters. All the experimental procedures performed on this isolate are the same as the ones for the isolates BEAP0338 and BEAP0335; this strain is deposited in the Roscoff Culture Collection, Roscoff Biological Station, National Center of Scientific Research, Sorbonne University, Roscoff, under ID 11331. The SEM stub is deposited in the Marine Biological Reference Collection of the Institute of Marine Science, Spanish Academy of Sciences, Barcelona, under ID XXX (will be provided prior to publication) (Santos-Bethencourt et al., 2023). The untrimmed, pairwise alignment of SSU rDNA sequences from HFCC157 (*P. azorensis*) and BEAP0342 produced in EMBOSS Needle tool (Madeira et al., 2024) is provided below.

668

```
BEAP0342          1 TGTCTAAGTGTAAATAAGTCTACACTGTGAAACTGCGAATGGCTCATTAA     50
                           |||| ||||||||||||||||||||||||||||||||||||||
HFCC157           1 -------GTGT-AATAAGTCTACACTGTGAAACTGCGAATGGCTCATTAA     42

BEAP0342         51 ATCAGTTATCGTTTATTTGATGGTCACTTACTACATGGATAACCGTAGTA    100
                    ||||||||||||||||||||||||||||||||||||||||||||||||||
HFCC157          43 ATCAGTTATCGTTTATTTGATGGTCACTTACTACATGGATAACCGTAGTA     92

BEAP0342        101 ATTCTAGAGCTAATACATGCATCAAGCCCCGACTTCGGAAGGGGTGTATT    150
                    ||||||||||||||||||||||||||||||||||||||||||||||||||
HFCC157          93 ATTCTAGAGCTAATACATGCATCAAGCCCCGACTTCGGAAGGGGTGTATT    142

BEAP0342        151 TATTAGATTCAAAACCAACCCTGGCAACAGGAACCTTGGTGATTCATAAT    200
                    ||||||||||||||||||||||||||||||||||||||||||||||||||
HFCC157         143 TATTAGATTCAAAACCAACCCTGGCAACAGGAACCTTGGTGATTCATAAT    192

BEAP0342        201 AACTTTTCGAACCGCATGGCCTTGAGCTGGCGGTGATTCATTCAAATTTC    250
                    ||||||||||||||||||||||||||||||||||||||||||||||||||
HFCC157         193 AACTTTTCGAACCGCATGGCCTTGAGCTGGCGGTGATTCATTCAAATTTC    242

BEAP0342        251 TGCCCTATCAACTTTCGATGGTAGGATAGAGGCCTACCATGGTTTTAACG    300
                    ||||||||||||||||||||||||||||||||||||||||||||||||||
HFCC157         243 TGCCCTATCAACTTTCGATGGTAGGATAGAGGCCTACCATGGTTTTAACG    292

BEAP0342        301 GGTGGCGGAGAATTAGGGTTCGATTCCGGAGAGGGAGCCTGAGAGACGGC    350
                    ||||.|||||||||||||||||||||||||||||||||||||||||||||
HFCC157         293 GGTGACGGAGAATTAGGGTTCGATTCCGGAGAGGGAGCCTGAGAGACGGC    342

BEAP0342        351 TACCACATCCAAGGAAGGCAGCAGGCGCGCAAATTACCCAATCCCAATAC    400
                    ||||||||||||||||||||||||||||||||||||||||||||||||||
HFCC157         343 TACCACATCCAAGGAAGGCAGCAGGCGCGCAAATTACCCAATCCCAATAC    392

BEAP0342        401 GGGGAGGTAGTGACAATAAATAACAATACCGGGCTCTCAGAGTCTGGTAA    450
                    ||||||||||||||||||||||||||||||||||||||||||||||||||
HFCC157         393 GGGGAGGTAGTGACAATAAATAACAATACCGGGCTCTCAGAGTCTGGTAA    442

BEAP0342        451 TTGGAATGAGAACAATTTAAATCCCTTAACGAGGATCAATTAGAGGGCAA    500
                    ||||||||||||||||||||||||||||||||||||||||||||||||||
HFCC157         443 TTGGAATGAGAACAATTTAAATCCCTTAACGAGGATCAATTAGAGGGCAA    492

BEAP0342        501 GTCTGGTGCCAGCAGCCGCGGTAATTCCAGCTCTAATAGCGTATATTAAA    550
                    ||||||||||||||||||||||||||||||||||||||||||||||||||
HFCC157         493 GTCTGGTGCCAGCAGCCGCGGTAATTCCAGCTCTAATAGCGTATATTAAA    542

BEAP0342        551 GTTGTTGCAGTTAAAAAGCTCGTAGTCGGATCTTGGGGTCGGGCAGGCTG    600
                    ||||||||||||||||||||||||||||||||||||||||||||||||||
HFCC157         543 GTTGTTGCAGTTAAAAAGCTCGTAGTCGGATCTTGGGGTCGGGCAGGCTG    592

BEAP0342        601 TCGGCCTAGTGTCGGACGGTCTTTTCGGCTCTTTCTTTCTGGGGA-----    645
                    |||||||||||||||||||||||||||||||||||||||||||||
HFCC157         593 TCGGCCTAGTGTCGGACGGTCTTTTCGGCTCTTTCTTTCTGGGGACTGCG    642

BEAP0342        646 -CTGCGTCTCTTTAATTAGGGGCGTATGGACGCAGATCGTTTACTTTGAA    694
                     |||||||||||||||||||||||||||||||||||||||||||||||||
HFCC157         643 TCTGCGTCTCTTTAATTAGGGGCGTATGGACGCAGATCGTTTACTTTGAA    692

BEAP0342        695 AAAATTAGAGTGTTCAAAGCAGGCTATCGCTTGAATACATTAGCATGGAA    744
                    ||||||||||||||||||||||||||||||||||||||||||||||||||
HFCC157         693 AAAATTAGAGTGTTCAAAGCAGGCTATCGCTTGAATACATTAGCATGGAA    742

BEAP0342        745 TAATGGAATAGGACTGCGGTTCTATTTTGTTGGTTTCTAGGACCGAAGTA    794
                    ||||||||||||||||||||||||||||||||||||||||||||||||||
HFCC157         743 TAATGGAATAGGACTGCGGTTCTATTTTGTTGGTTTCTAGGACCGAAGTA    792

BEAP0342        795 ATGATTAATAGGGATAGTTGGGGGCCGTTTATATTCCGTTGTCAGAGGTGA    844
                    ||||||||||||||||||||||||||||||||||||||||||||||||||
HFCC157         793 ATGATTAATAGGGATAGTTGGGGGCCGTTTATATTCCGTTGTCAGAGGTGA    842

BEAP0342        845 AATTCTTGGATTTACGGAAGATAAACTTCTGCGAAAGCATTCGGCAAGGA    894
                    ||||||||||||||||||||||||||||||||||||||||||||||||||
HFCC157         843 AATTCTTGGATTTACGGAAGATAAACTTCTGCGAAAGCATTCGGCAAGGA    892

BEAP0342        895 TGTTTTCATTGATCAAGAACGAAAGTTAGGGGATCGAAGACGATCAGATA    944
                    ||||||||||||||||||||||||||||||||||||||||||||||||||
HFCC157         893 TGTTTTCATTGATCAAGAACGAAAGTTAGGGGATCGAAGACGATCAGATA    942
```

669

```
BEAP0342         945 CCGTCGTAGTCTTAACCATAAACTATGCCGACTAGGGATCAGTGGACGTT    994
                     ||||||||||||||||||||||||||||||||||||||||||||||||||
HFCC157          943 CCGTCGTAGTCTTAACCATAAACTATGCCGACTAGGGATCAGTGGACGTT    992

BEAP0342         995 GTTTTACGACTTCATTGGCACCTTGTGAGAAATCAAAGTTTTTGGGTTCC   1044
                     ||||||||||||||||||||||||||||||||||||||||||||||||||
HFCC157          993 GTTTTACGACTTCATTGGCACCTTGTGAGAAATCAAAGTTTTTGGGTTCC   1042

BEAP0342        1045 GGGGGGAGTATGGTCGCAAGGCTGAAACTTAAAGGAATTGACGGAAGGGC   1094
                     ||||||||||||||||||||||||||||||||||||||||||||||||||
HFCC157         1043 GGGGGGAGTATGGTCGCAAGGCTGAAACTTAAAGGAATTGACGGAAGGGC   1092

BEAP0342        1095 ACCACCAGGAGTGGAGCCTGCGGCTTAATTTGACTCAACACGGGGAAACT   1144
                     ||||||||||||||||||||||||||||||||||||||||||||||||||
HFCC157         1093 ACCACCAGGAGTGGAGCCTGCGGCTTAATTTGACTCAACACGGGGAAACT   1142

BEAP0342        1145 TACCAGGTCCAGACATAGTAAGGATTGACAGATTGAAAGCTCTTTCTTGA   1194
                     ||||||||||||||||||||||||||||||||||||||||||||||||||
HFCC157         1143 TACCAGGTCCAGACATAGTAAGGATTGACAGATTGAAAGCTCTTTCTTGA   1192

BEAP0342        1195 TTCTATGGGTGGTGGTGCATGGCCGTTCTTAGTTGGTGGAGTGATTTGTC   1244
                     ||||||||||||||||||||||||||||||||||||||||||||||||||
HFCC157         1193 TTCTATGGGTGGTGGTGCATGGCCGTTCTTAGTTGGTGGAGTGATTTGTC   1242

BEAP0342        1245 TGGTTAATTCCGTTAACGAACGAGACCTCAGCCTACTAAATAGTCACGCG   1294
                     ||||||||||||||||||||||||||||||||||||||||||||||||||
HFCC157         1243 TGGTTAATTCCGTTAACGAACGAGACCTCAGCCTACTAAATAGTCACGCG   1292

BEAP0342        1295 AAGTTTTACTTCGTGGCCGGCTTCTTAGAGGGACTATTTGTGTTTAACGA   1344
                     ||||||||||||||||||||||||||||||||||||||||||||||||||
HFCC157         1293 AAGTTTTACTTCGTGGCCGGCTTCTTAGAGGGACTATTTGTGTTTAACGA   1342

BEAP0342        1345 ATGGAAGTTTGAGGCAATAACAGGTCTGTGATGCCCTTAGATGTTCTGGG   1394
                     ||||||||||||||||||||||||||||||||||||||||||||||||||
HFCC157         1343 ATGGAAGTTTGAGGCAATAACAGGTCTGTGATGCCCTTAGATGTTCTGGG   1392

BEAP0342        1395 CCGCACGCGCGCTACACTGATGAATTCAACGAGCTCACAACCTTGACCGA   1444
                     ||||||||||||||||||||||||||||||||||||||||||||||||||
HFCC157         1393 CCGCACGCGCGCTACACTGATGAATTCAACGAGCTCACAACCTTGACCGA   1442

BEAP0342        1445 AAGGCCCGGGTAAACTCCGAAATTTCATCGTGATGGGGATAGACTATTGT   1494
                     ||||||||||||||||||||||||||||||||||||||||||||||||||
HFCC157         1443 AAGGCCCGGGTAAACTCCGAAATTTCATCGTGATGGGGATAGACTATTGT   1492

BEAP0342        1495 AATTATTAGTCTTCAACGAGGAATTCCTAGTAAGCGCGATTCATCAGATC   1544
                     ||||||||||||||||||||||||||||||||||||||||||||||||||
HFCC157         1493 AATTATTAGTCTTCAACGAGGAATTCCTAGTAAGCGCGATTCATCAGATC   1542

BEAP0342        1545 GCGTTGATTACGTCCCTGCCCTTTGTACACACCGCCCGTCGCTACTACCG   1594
                     ||||||||||||||||||||||||||||||||||||||||||||||||||
HFCC157         1543 GCGTTGATTACGTCCCTGCCCTTTGTACACACCGCCCGTCGCTACTACCG   1592

BEAP0342        1595 ATTGAATGGCTTAGTGAGGCTCCCGGACCGACGATTGGTGGCTTCACGGC   1644
                     ||||||||||||||||||||||||||||||||||||||||||||||||||
HFCC157         1593 ATTGAATGGCTTAGTGAGGCTCCCGGACCGACGATTGGTGGCTTCACGGC   1642

BEAP0342        1645 TGCTAGTTGTTGGAAAGTTAGACAAACTTGGTCATTTAGAGGAAGTAAAA   1694
                     ||||||||
HFCC157         1643 TGCTAGTT------------------------------------------   1650

BEAP0342        1695 GTCGTAACAAGGTTTCC   1711

HFCC157         1651 -----------------   1650
```

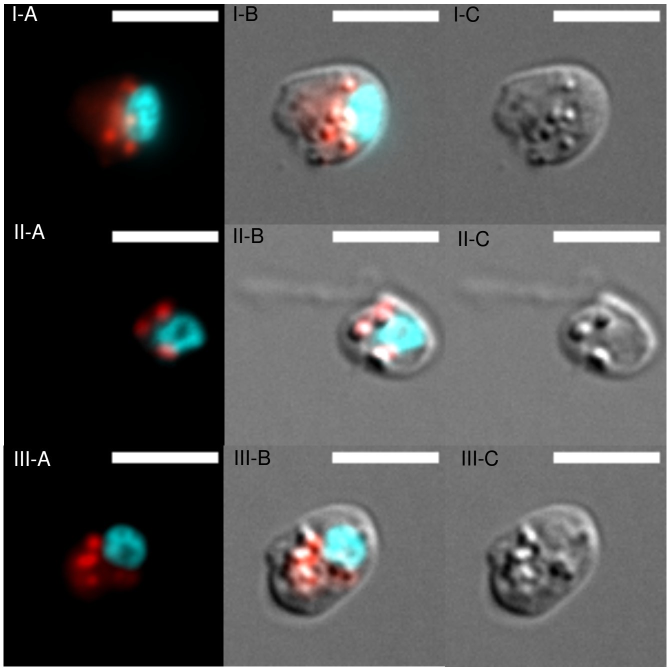


Figure S2: **Lipid bodies and nuclei of BEAP0340 stained with Nile Red and Hoechst.** Three different cells are presented under number I, II and III; Red - Nile Red staining, blue - Hoechst, grey - DIC. A - Nile Red + Hoechst, B - Nile Red + Hoechst + DIC, C - DIC. Scale bar 5 µm. The co-localization of DIC-contrasted globules and lipid-binding dye is visible. Due to cytotoxicity of Nile Red with DMSO, cells lost flagella and discharged ejectosomes; however, the presence of an intact nucleus and overall preservation of cell shape indicate that cell death did not start yet (all images were acquired <10min after adding Nile Red). Live cells were stained for 15 minutes with Hoechst diluted 1:1000, and treated with Nile Red solution in DMSO (final concentration of Nile Red 50 ng/µl). The images were acquired with an Axio Observer.Z1 inverted microscope a DIC Plan-Apochromat 63x Oil immersion objective (NA=1.4), an HDCamC13440-20CU Hamamatsu camera and ZEN image acquisition software. Images were subsequently reconstructed using Fiji (Schindelin et al., 2012).

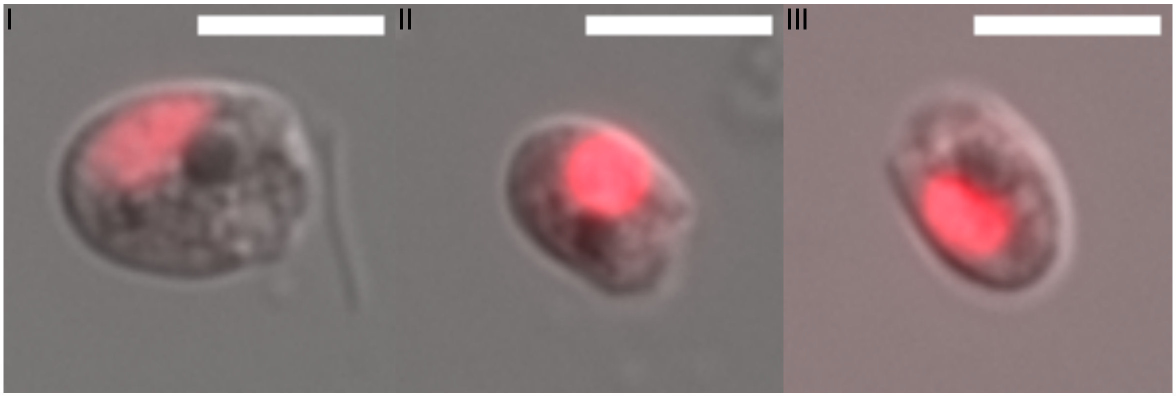


Figure S3: **Nuclei of BEAP0338 stained with DRAQ5.** Three different cells are presented under number I, II and III; Red - DRAQ5, grey - DIC. The dark spherical structure is visible in DIC image, while the nucleus stained with DRAQ5 is a different structure, shifted to the posterior side of the cell. Scale bar 5 µm. Live cells were stained for 15 minutes with DRAQ5 (final concentration of DRAQ5 0.0455mM). The images were acquired with an Axio Observer.Z1 inverted microscope a DIC Plan-Apochromat 63x Oil immersion objective (NA=1.4), an HDCamC13440-20CU Hamamatsu camera and ZEN image acquisition software. Images were subsequently reconstructed using Fiji (Schindelin et al., 2012).

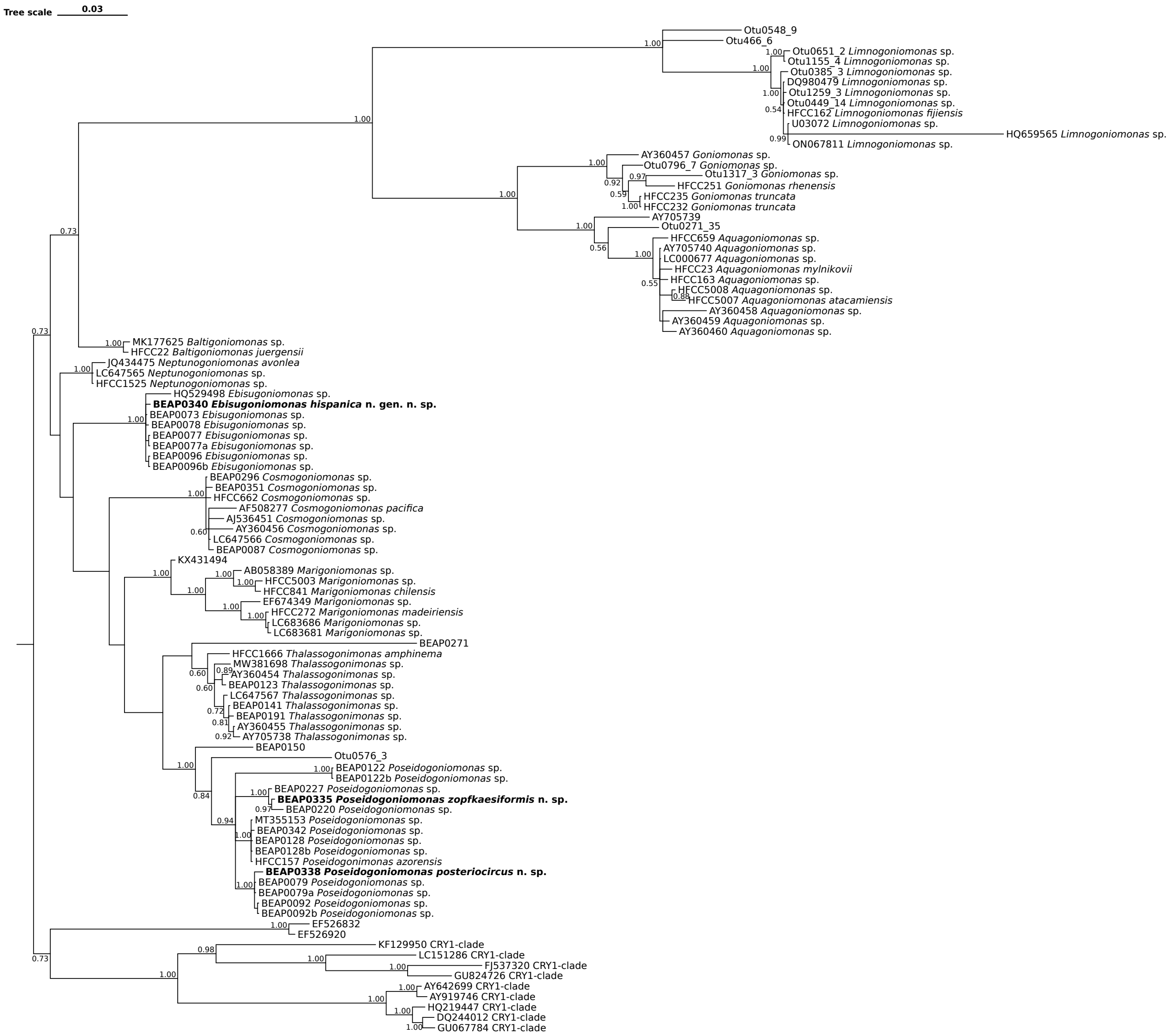


Figure S4: **Bayesian phylogeny of goniomonads.** The values at the nodes represent the posterior probability. The scale represents substitutions per site.

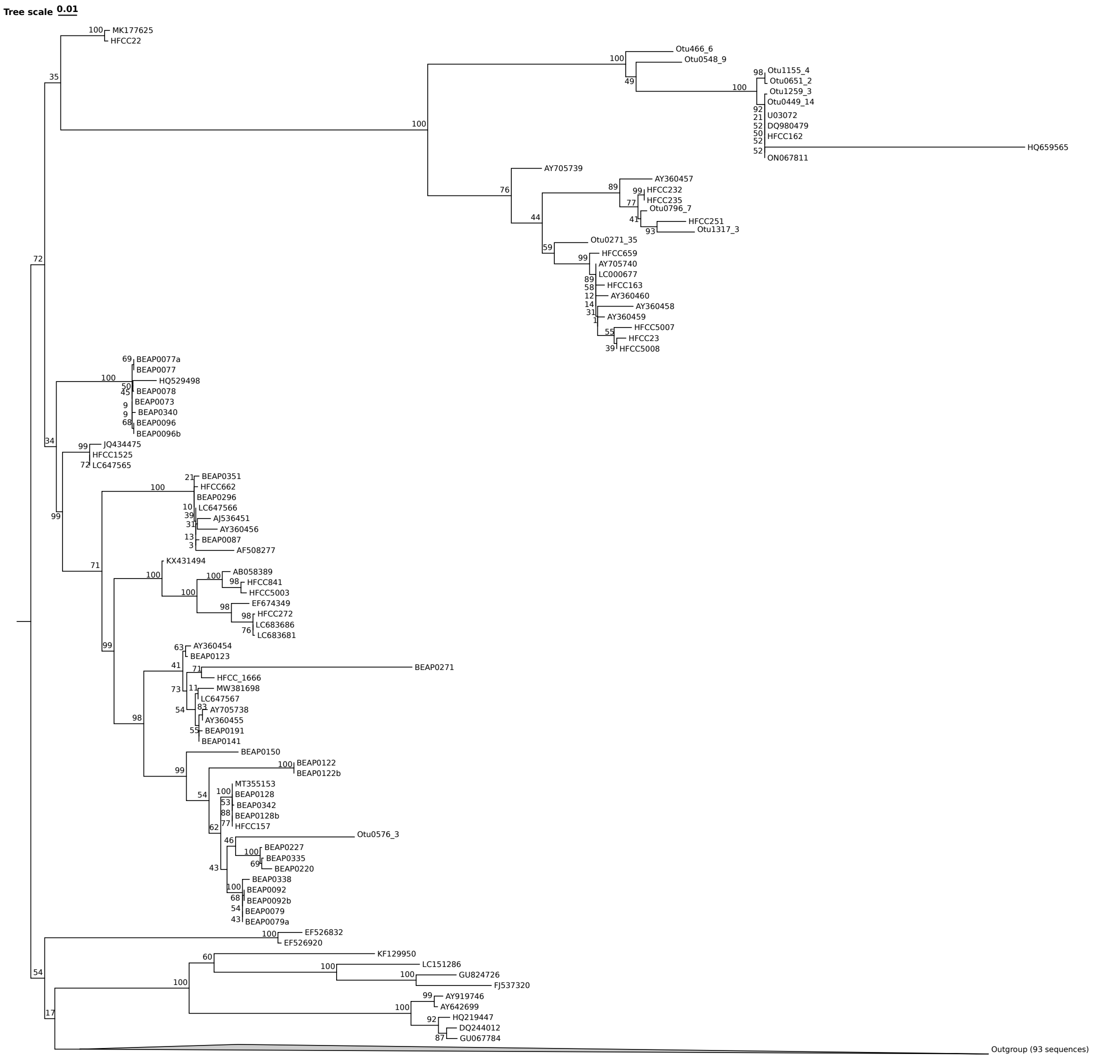


Figure S5: **Maximum Likelihood phylogeny of goniomonads with expanded taxa selection for an outgroup.** The values at the nodes represent bootstrap support. The closest lineage from the outgroup, which contains environmental DNA sequences EF526920 and EF526832 and CRY-1 lineages were expanded in this representation. The larger outgroup contains 93 sequences of representatives from all major Cryptista lineages. The scale represents substitutions per site. The process of obtaining this phylogeny is the same as the one described in Materials and Methods.

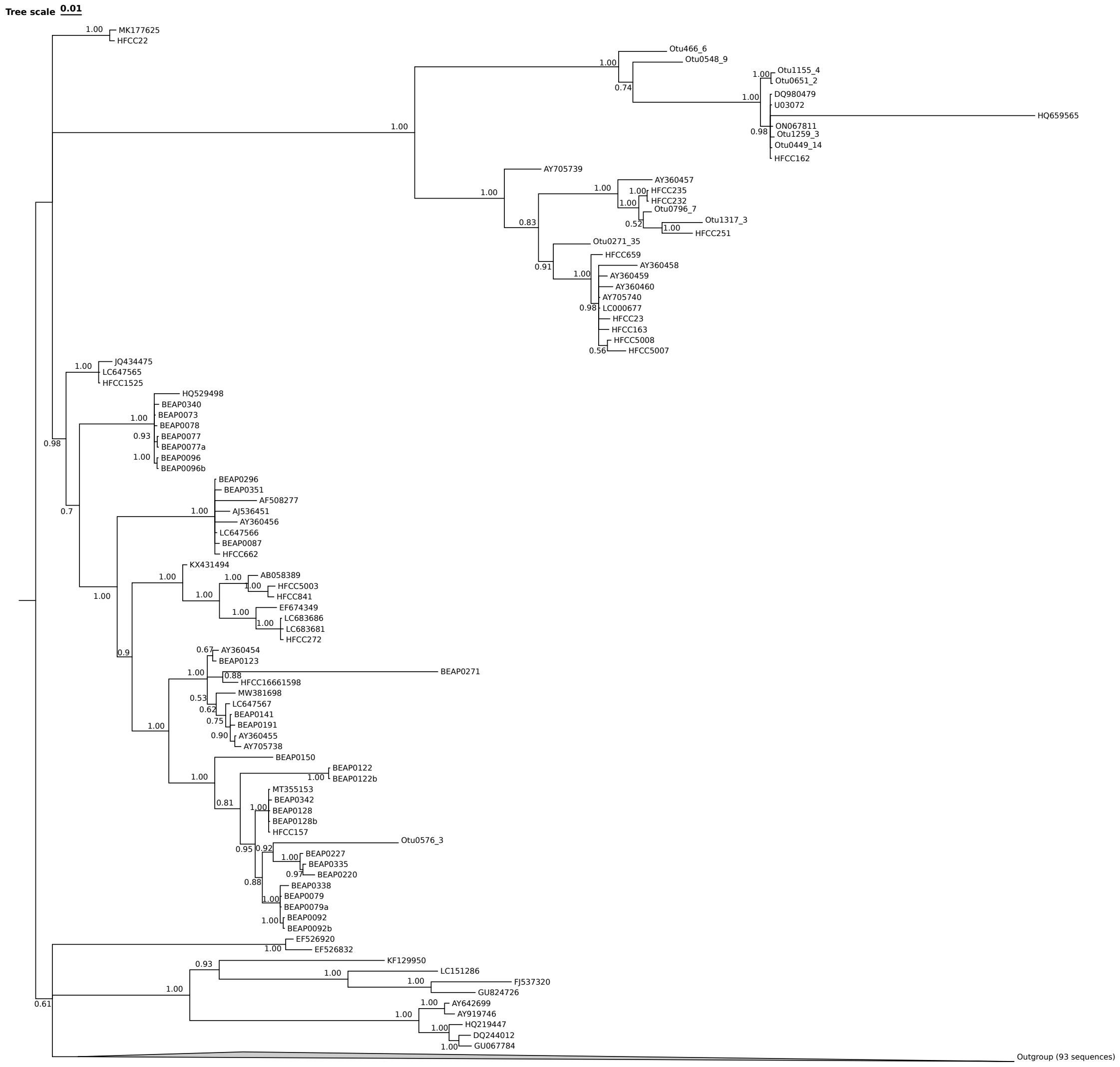


Figure S6: **Bayesian phylogeny of goniomonads with expanded taxa selection for an outgroup.** The values at the nodes represent posterior probability. The closest lineage from the outgroup, which contains environmental DNA sequences EF526920 and EF526832 and CRY-1 lineages were expanded in this representation. The larger outgroup contains 93 sequences of representatives from all major Cryptista lineages. The scale represents substitutions per site. The process of obtaining this phylogeny is the same as the one described in Materials and Methods.

:

:

Movie S1: **Predominating movement pattern of BEAP0340.** https://figshare.com/articles/media/Goniomonad_species_imaging_data/25988158?file=52189400 The images were acquired with an Axio Observer.Z1 inverted microscope a DIC Plan-Apochromat 63x Oil immersion objective (NA=1.4), an HDCamC13440-20CU Hamamatsu camera and ZEN image acquisition software. Images were subsequently reconstructed using Fiji (Schindelin et al., 2012).

Movie S2: **Predominating movement pattern of BEAP0338.** https://figshare.com/articles/media/Goniomonad_species_imaging_data/25988158?file=52189397 The images were acquired with an Axio Observer.Z1 inverted microscope a DIC Plan-Apochromat 63x Oil immersion objective (NA=1.4), an HDCamC13440-20CU Hamamatsu camera and ZEN image acquisition software. Images were subsequently reconstructed using Fiji (Schindelin et al., 2012).

Movie S3: **Predominating movement pattern of BEAP0335.** https://figshare.com/articles/media/Goniomonad_species_imaging_data/25988158?file=52189391 The images were acquired with an Axio Observer.Z1 inverted microscope a DIC Plan-Apochromat 63x Oil immersion objective (NA=1.4), an HDCamC13440-20CU Hamamatsu camera and ZEN image acquisition software. Images were subsequently reconstructed using Fiji (Schindelin et al., 2012).

| **RaXML** | BEAP0340 | BEAP0073 | BEAP0077 | BEAP0096 | HQ529498 |
|---|---|---|---|---|---|
| BEAP0340 | – | 0,16% | 0,24% | 0,24% | 1,35% |
| BEAP0073 | 0,16% | – | 0,08% | 0,08% | 1,19% |
| BEAP0077 | 0,24% | 0,08% | – | 0,16% | 1,27% |
| BEAP0096 | 0,24% | 0,08% | 0,16% | – | 1,27% |
| HQ529498 | 1,35% | 1,19% | 1,27% | 1,27% | – |
| **MrBayes** | BEAP0340 | BEAP0073 | BEAP0077 | BEAP0096 | HQ529498 |
| BEAP0340 | – | 0,25% | 0,37% | 0,37% | 1,28% |
| BEAP0073 | 0,25% | – | 0,23% | 0,23% | 1,13% |
| BEAP0077 | 0,37% | 0,23% | – | 0,35% | 1,26% |
| BEAP0096 | 0,37% | 0,23% | 0,35% | – | 1,26% |
| HQ529498 | 1,28% | 1,13% | 1,26% | 1,26% | – |

Table S1: **Pairwise cophenetic distances for sequences that belong to *Ebisugoniomonas* clade.** Distances calculated from both ML("RaXML") and BI("MrBayes") phylogenetic trees are represented. If several sequences originating from the same isolate had a distance <0.01%, only one sequence was chosen.

| **RaXML** | BEAP0079 | BEAP0128 | BEAP0220 | BEAP0335 | BEAP0338 | BEAP0342 | BEAP0092 | BEAP0122 | BEAP0150 | BEAP0227 | HFCC157 (P. azorensis) | MT355153 |
|---|---|---|---|---|---|---|---|---|---|---|---|---|
| BEAP0079 | – | 1,66% | 3,00% | 2,59% | 0,34% | 1,74% | 0,08% | 5,68% | 5,35% | 2,51% | 1,66% | 1,66% |
| BEAP0128 | 1,66% | – | 2,98% | 2,57% | 2,00% | 0,08% | 1,74% | 5,23% | 5,33% | 2,49% | <0.01% | <0.01% |
| BEAP0220 | 3,00% | 2,98% | – | 0,58% | 3,35% | 3,06% | 3,09% | 7,00% | 6,52% | 0,66% | 2,98% | 2,98% |
| BEAP0335 | 2,59% | 2,57% | 0,58% | – | 2,93% | 2,65% | 2,67% | 6,59% | 6,10% | 0,25% | 2,57% | 2,57% |
| BEAP0338 | 0,34% | 2,00% | 3,35% | 2,93% | – | 2,08% | 0,42% | 6,02% | 5,69% | 2,85% | 2,00% | 2,00% |
| BEAP0342 | 1,74% | 0,08% | 3,06% | 2,65% | 2,08% | – | 1,82% | 5,31% | 5,41% | 2,57% | 0,08% | 0,08% |
| BEAP0092 | 0,08% | 1,74% | 3,09% | 2,67% | 0,42% | 1,82% | – | 5,76% | 5,43% | 2,59% | 1,74% | 1,74% |
| BEAP0122 | 5,68% | 5,23% | 7,00% | 6,59% | 6,02% | 5,31% | 5,76% | – | 9,34% | 6,50% | 5,23% | 5,23% |
| BEAP0150 | 5,35% | 5,33% | 6,52% | 6,10% | 5,69% | 5,41% | 5,43% | 9,34% | – | 6,02% | 5,33% | 5,33% |
| BEAP0227 | 2,51% | 2,49% | 0,66% | 0,25% | 2,85% | 2,57% | 2,59% | 6,50% | 6,02% | – | 2,49% | 2,49% |
| HFCC157 (P. azorensis) | 1,66% | <0.01% | 2,98% | 2,57% | 2,00% | 0,08% | 1,74% | 5,23% | 5,33% | 2,49% | – | <0.01% |
| MT355153 | 1,66% | <0.01% | 2,98% | 2,57% | 2,00% | 0,08% | 1,74% | 5,23% | 5,33% | 2,49% | <0.01% | – |
| **MrBayes** | BEAP0079 | BEAP0128 | BEAP0220 | BEAP0335 | BEAP0338 | BEAP0342 | BEAP0092 | BEAP0122 | BEAP0150 | BEAP0227 | HFCC157 (P. azorensis) | MT355153 |
| BEAP0079 | – | 1,60% | 2,93% | 2,56% | 0,41% | 1,68% | 0,24% | 5,08% | 5,07% | 2,43% | 1,60% | 1,60% |
| BEAP0128 | 1,60% | – | 2,78% | 2,41% | 1,91% | 0,18% | 1,73% | 4,93% | 4,92% | 2,28% | 0,10% | 0,10% |
| BEAP0220 | 2,93% | 2,78% | – | 0,63% | 3,24% | 2,86% | 3,06% | 6,26% | 6,25% | 0,76% | 2,78% | 2,78% |
| BEAP0335 | 2,56% | 2,41% | 0,63% | – | 2,87% | 2,50% | 2,69% | 5,89% | 5,89% | 0,40% | 2,42% | 2,41% |
| BEAP0338 | 0,41% | 1,91% | 3,24% | 2,87% | – | 1,99% | 0,54% | 5,38% | 5,38% | 2,74% | 1,91% | 1,90% |
| BEAP0342 | 1,68% | 0,18% | 2,86% | 2,50% | 1,99% | – | 1,81% | 5,01% | 5,01% | 2,37% | 0,19% | 0,18% |
| BEAP0092 | 0,24% | 1,73% | 3,06% | 2,69% | 0,54% | 1,81% | – | 5,20% | 5,20% | 2,56% | 1,73% | 1,73% |
| BEAP0122 | 5,08% | 4,93% | 6,26% | 5,89% | 5,38% | 5,01% | 5,20% | – | 8,40% | 5,76% | 4,93% | 4,92% |
| BEAP0150 | 5,07% | 4,92% | 6,25% | 5,89% | 5,38% | 5,01% | 5,20% | 8,40% | – | 5,75% | 4,93% | 4,92% |
| BEAP0227 | 2,43% | 2,28% | 0,76% | 0,40% | 2,74% | 2,37% | 2,56% | 5,76% | 5,75% | – | 2,29% | 2,28% |
| HFCC157 | 1,60% | 0,10% | 2,78% | 2,42% | 1,91% | 0,19% | 1,73% | 4,93% | 4,93% | 2,29% | – | 0,10% |
| MT355153 | 1,60% | 0,10% | 2,78% | 2,41% | 1,90% | 0,18% | 1,73% | 4,92% | 4,92% | 2,28% | 0,10% | – |

Table S2: **Pairwise cophenetic distances for sequences that belong to *Poseidogoniomonas* clade and isolate BEAP0150.** Distances calculated from both ML("RaXML") and BI("MrBayes") phylogenetic trees are represented. Otu0576_3 was omitted; if several sequences originating from the same isolate had a distance <0.01%, only one sequence was chosen.
